# Stepwise assembly of virulence-associated traits in the intracellular pathogen *Coxiella burnetii*

**DOI:** 10.64898/2026.05.14.724635

**Authors:** Amanda E. Brenner, Rahul Raghavan

## Abstract

*Coxiella burnetii* is the only member of the order Legionellales known to primarily infect vertebrates. The Q fever pathogen is also unusual in that it replicates within an acidified phagolysosome-like vacuole. The evolutionary origins of the virulence determinants underlying this lifestyle remain unclear. More broadly, little is known about how virulence-related traits arise in specialized intracellular lineages, where access to foreign-origin DNA may be more episodic. To address this question, we used Legionellales-wide comparative phylogenomics to reconstruct the gain and loss of traits affecting host interaction, immune evasion, intracellular survival, and metabolism. We found that many virulence-associated traits in *C. burnetii* predate the modern pathogen and were assembled stepwise in ancestors that likely occupied niches distinct from the acidified vacuolar niche of modern *C. burnetii*. The common ancestor shared with soft-tick *Coxiella* endosymbionts likely encoded most *C. burnetii* type IVB secretion system effectors, indicating that much of the host-manipulation repertoire in *C. burnetii* was already present before the emergence of the modern pathogen. Distinctive lipopolysaccharide features associated with immune evasion also appear to have accumulated progressively within the *Coxiella* lineage, including genes implicated in synthesis of virenose, a unique O-antigen sugar critical for *C. burnetii* virulence. Traits likely to support replication in the acidic *Coxiella*-containing vacuole likewise accumulated gradually, with generalized stress-tolerance functions predating acquisition of an Mrp cation/proton antiporter that may have further supported pH homeostasis. Additional changes in sugar transport and catabolism, glycolytic control, and respiratory metabolism likely enhanced metabolic flexibility and access to diverse substrates in this nutrient-rich niche. Together, these findings support a model in which vertebrate pathogenicity in *C. burnetii* emerged through stepwise remodeling of an ancestral host-associated lineage and provide a framework for understanding how virulence-related traits evolve in specialized intracellular pathogens.

**AUTHOR SUMMARY:** *Coxiella burnetii* is the bacterium that causes Q fever, a disease that can spread from animals to humans. Unlike its close relatives, *C. burnetii* primarily infects vertebrates and grows inside an acidic compartment within host cells. New bacterial pathogens often evolve by gaining genes from other bacteria, but how virulence evolves in lineages that grow only inside host cells, where opportunities to gain new genes may be infrequent, remains unclear. We wanted to understand how *C. burnetii* evolved the traits needed for its distinctive intracellular lifestyle. By comparing its genome to those of related bacteria across the order Legionellales, we found that features involved in host manipulation, immune evasion, acid tolerance, and nutrient use appeared at different times in its ancestry rather than being acquired all at once by the modern pathogen. Our findings suggest that specialized intracellular pathogens can emerge through gradual changes in ancestral host-associated lineages, including gene acquisition, gene loss, retention of older traits, and repurposing of existing functions.

## INTRODUCTION

The order Legionellales contains several intracellular bacteria that infect eukaryotic hosts, but *Coxiella burnetii*, the causative agent of Q fever, is the only known member that primarily infects vertebrates [1–8]. This unusual host range is coupled to a distinctive intracellular niche: *C. burnetii* replicates within a large *Coxiella*-containing vacuole (CCV) that matures through the canonical endocytic pathway and acquires phagolysosomal characteristics [9–11]. As the vacuole acidifies to approximately pH 4.75, dormant small cell variants transition into metabolically active large cell variants and assemble the Dot/Icm type IVB secretion system (T4BSS), which is essential for intracellular replication [12–16]. The mature CCV is therefore both hostile and nutrient rich, exposing *C. burnetii* to low pH, lysosomal hydrolases, reactive oxygen species, and osmotic stress while also providing access to diverse host-derived substrates [17–20]. Understanding how *C. burnetii* evolved the ability to survive and replicate in this compartment is central to uncovering the origin of vertebrate pathogenicity in the genus.

Several traits are known to be critical to *C. burnetii* pathogenesis. First, the Dot/Icm T4BSS delivers effector proteins that modulate host trafficking pathways and promote fusion of the CCV with endosomes, lysosomes, and autophagosomes, thereby enabling vacuole expansion and bacterial growth [9,10,13,21]. Second, *C. burnetii* LPS contributes to immune evasion, and loss of the full-length O-antigen prevents productive infection in immunocompetent vertebrates [22–24]. This O-antigen contains the unusual sugars virenose and dihydrohydroxystreptose, which are rarely found in other bacteria and are essential for normal O-antigen extension [23, 25–27]. Third, *C. burnetii* is more metabolically flexible relative to other Legionellales and can efficiently use both amino acids and sugars as carbon and energy sources, a feature that may be advantageous in the chemically diverse environment of the CCV [28–30]. Together, these features suggest that vertebrate pathogenicity in *C. burnetii* depends on host-cell manipulation, immune evasion, adaptation to acidic intracellular conditions, and metabolic versatility.

The evolutionary origins of these traits remain incompletely resolved. Earlier work proposed that *C. burnetii* arose from a maternally inherited tick endosymbiont that acquired the factors needed to infect vertebrate cells [31]. More recent analyses instead indicate that *C. burnetii* and closely related *Coxiella* endosymbionts (CEs) descended from pathogenic ancestors and that the common ancestor of *C. burnetii* and soft-tick endosymbionts likely already possessed T4BSS and the capacity to infect macrophage-like cells [32,33]. This shift in perspective reframes the problem from the origin of pathogenicity itself to the origin of the more specialized traits that enabled *C. burnetii* to exploit vertebrate hosts and thrive in an acidic intracellular vacuole.

To address this question, we used comparative phylogenomics across Legionellales, together with a newly assembled genome of the CE in *Ornithodoros peruvianus*, to trace the evolution of traits associated with vertebrate pathogenicity in *C. burnetii* (**Figure 1**). This genome is especially informative for reconstructing *C. burnetii* evolution because *Ornithodoros*-associated CEs are the closest sampled relatives of the pathogen and preserve a comparatively broad set of orthologous and pseudogenized genes shared with *C. burnetii* [34] (**Table 1**). We focus particularly on systems likely to influence host interaction, immune evasion, survival in acidic intracellular conditions, and exploitation of host-derived nutrients. Our results indicate that much of *C. burnetii*’s virulence-associated machinery predates the modern pathogen, and that key features linked to vertebrate adaptation, including aspects of LPS structure, acid tolerance, and metabolic flexibility, were assembled more gradually. Together, these findings support a model in which modern *C. burnetii* emerged through stepwise remodeling of an ancestral host-associated lineage rather than through a recent transition from a benign tick symbiont.

**Figure 1.**
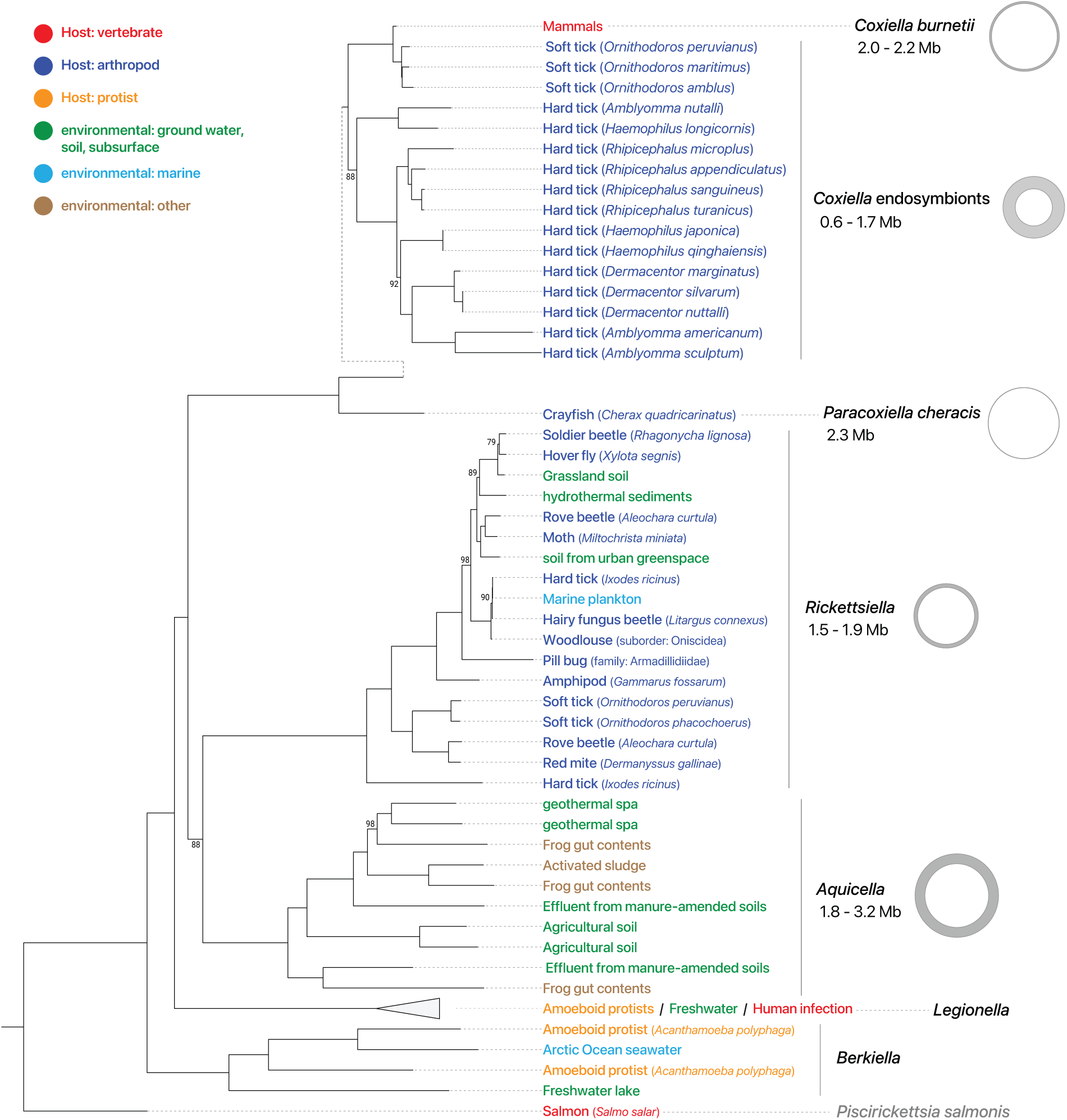
Legionellales members are found in diverse hosts or environments and have varied genome sizes. Tips of the tree are colored to indicate the host or environment from which they were collected. Circles represent the range of genome sizes within the genus. The *Legionella sp.* clade has been collapsed for brevity. All nodes have bootstrap values of 100 except for those with listed values. The *Coxiella* ML tree (top) was generated using 78 protein sequences (Table S2). The Legionellales ML tree (bottom) was generated using 152 protein sequences (Table S3).

**Table 1.** *Coxiella* genome characteristics.

| Taxa | Genome Size (Mb) | Protein Coding Genes | Pseudogenes | Orthogroups | Assembly Accession |
| --- | --- | --- | --- | --- | --- |
| *Coxiella burnetii* RSA 493 | 2.033 | 1,833 | 207 | 1758 | GCF_000007765.2 |
| CE *Ornithodoros peruvianus* | 1.672 | 1,250 | 638 | 1523 |  |
| CE *Ornithodoros amblus* | 1.561 | 1,049 | 534 | 1352 | GCF_019425495.1 |
| CE *Ornithodoros maritimus* | 1.671 | 1,225 | 622 | 1475 | GCF_907164965.1 |
| CE *Amblyomma nuttalli* | 1.003 | 668 | 16 | 657 | GCF_018107685.1 |
| CE *Haemaphysalis longicornis* | 0.987 | 716 | 26 | 718 | GCA_048491735.1 |
| CE *Rhipicephalus microplus* | 1.565 | 638 | 163 | 684 | GCF_002871095.1 |
| CE *Rhipicephalus appendiculatus* | 1.449 | 1,222 | 206 | 1190 | GCF_030643785.1 |
| CE *Rhipicephalus sanguineus* | 1.716 | 1,535 | 104 | 1272 | GCF_002804145.1 |
| CE *Rhipicephalus turanicus* | 1.734 | 1,626 | 151 | 1329 | GCF_001077715.1 |
| CE *Haemaphysalis japonica* | 0.878 | 679 | 16 | 666 | GCA_048491875.1 |
| CE *Haemaphysalis qinghaiensis* | 0.884 | 689 | 18 | 674 | GCA_048492375.1 |
| CE *Dermacentor marginatus* | 0.901 | 637 | 18 | 630 | GCF_907164955.1 |
| CE *Dermacentor silvarum* | 0.887 | 637 | 17 | 633 | GCA_048493135.1 |
| CE *Dermacentor nuttalli* | 0.888 | 641 | 18 | 639 | GCA_048494595.1 |
| CE *Amblyomma americanum* | 0.657 | 559 | 7 | 557 | GCF_000815025.1 |
| CE *Amblyomma americanum* | 0.657 | 559 | 7 | 557 | GCF_000815025.1 |
| CE *Amblyomma sculptum* | 0.623 | 526 | 9 | 528 | GCF_009883795.1 |

## RESULTS

### Ancient T4BSS machinery was paired with a more recently assembled *C. burnetii* effector repertoire

The Dot/Icm T4BSS is essential for virulence in Legionellales [12,35–37]. Analysis of T4BSS components revealed that most are broadly conserved throughout Legionellales, with the main exception being the CEs (**Figure 2**). In contrast to the T4BSS machinery, most effectors were found only in close relatives of *Coxiella*. Of the 28 *C. burnetii* effectors with established host interactions [21], 20 have orthologs in one or more *Ornithodoros* CEs and six have orthologs in *Paracoxiella* (**Table 2**). Few effectors could be traced beyond the common ancestor of *Coxiella* and *Paracoxiella*: seven lacked any orthologs elsewhere in the Legionellales, and six had no orthologs outside *C. burnetii*. These data suggest that many *C. burnetii* effectors arose relatively recently and are tailored to its particular host range and intracellular lifestyle.

**Figure 2.**
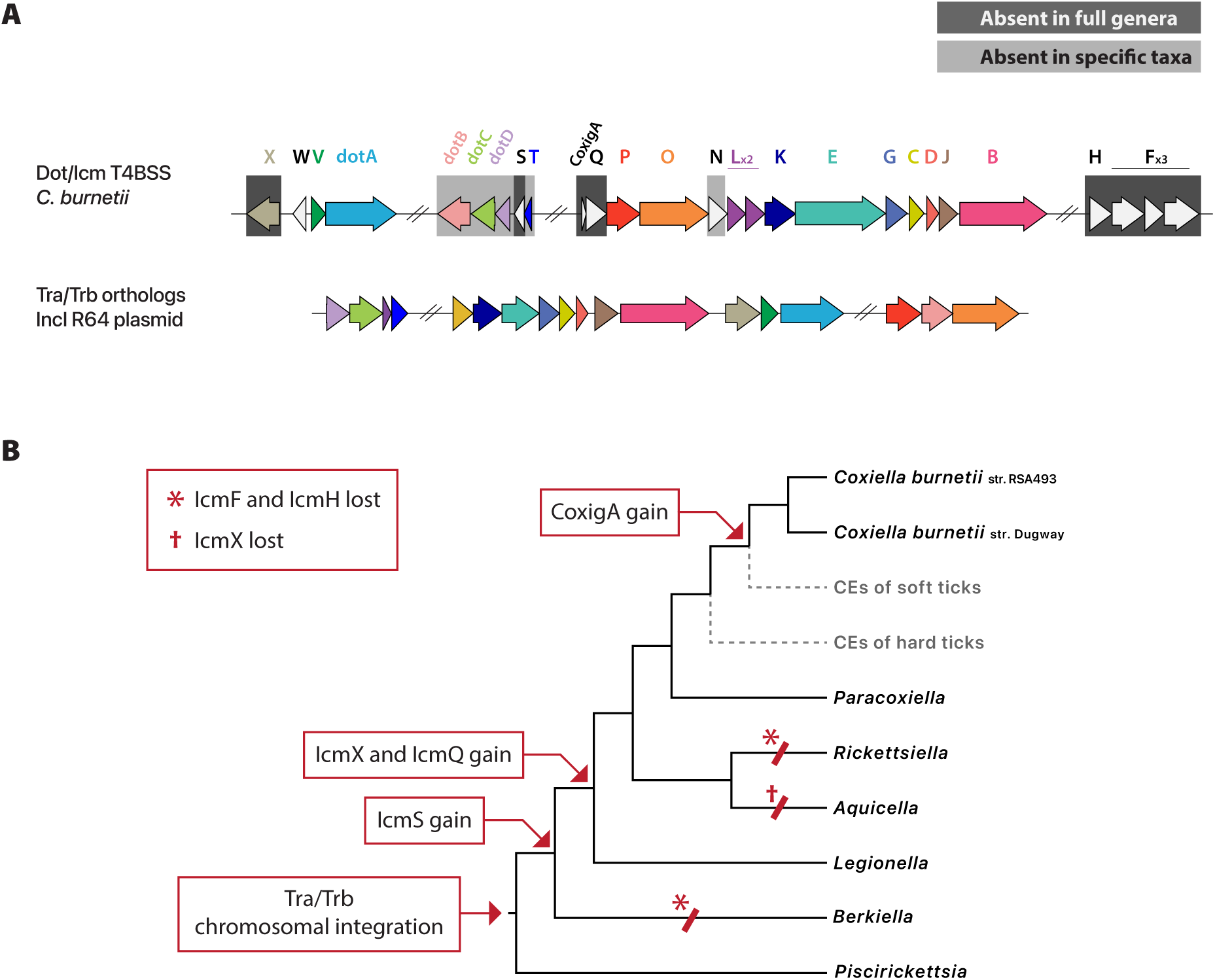
Most type IVB secretion system components are present throughout Legionellales. **(A**) Conservation of T4BSS genes present in *C. burnetii*: genes shown in color have homologs in Tra/Trb conjugation system (shown below), genes highlighted in grey boxes are absent in one or more Legionellales taxa, and genes without highlights have been maintained throughout Legionellales, excluding the CEs. (**B**) Summary of major gains (red arrows) and losses (red crosses, red asterisks) in T4BSS evolution. Dashed gray lines indicate branches where T4BSS has been lost or pseudogenized.

**Table 2.** Presence of *C. burnetii* effector orthologs in CEs or *Paracoxiella*.

| locus | Acronym | *Coxiella* Endosymbionts | *Paracoxiella cheracis* |
| --- | --- | --- | --- |
| CBU_0175 | CstK | ✓ | ✓ |
| CBU_0513 | CinF | ✓ | ✓ |
| CBU_0937 | MceB/CirC | ✓ | ✓ |
| CBU_1425 | MceC | ✓ | ✓ |
| CBU_1863 | CvpE | ✓ | ✓ |
| CBU_0021 | CvpB/Cig2 | ✓ |  |
| CBU_0041 | CirA/CoxCC1 | ✓ |  |
| CBU_0077 | MceA | ✓ |  |
| CBU_0122 | CvpM | ✓ |  |
| CBU_0388 | CetCb2 | ✓ |  |
| CBU_0447 | AnkF | ✓ |  |
| CBU_0635 | - | ✓ |  |
| CBU_0781 | AnkG | ✓ |  |
| CBU_1370 | CbEPF1 | ✓ |  |
| CBU_1387 | EmcA/Cem6 | ✓ |  |
| CBU_1751 | Cig57 | ✓ |  |
| CBU_2007 | Vice | ✓ |  |
| CBU_2013 | EmcB | ✓ |  |
| CBUD_0462 | CaeA | ✓ |  |
| CBUA0013 | CpeB |  | ✓ |
| CBU_0425 | CirB |  |  |
| CBU_0626 | CvpF |  |  |
| CBU_0665 | CvpA |  |  |
| CBU_1217 | NopA |  |  |
| CBU_1314 | coxCC6 |  |  |
| CBU_1531 | CaeB |  |  |
| CBU_1823 | IcaA |  |  |

Our analysis also indicates that the effector MceB (CirC, CBU_0937) and the hypothetical protein CBU_1699 arose through duplication of the LbtP siderophore porin, which is present throughout Legionellales (**Figure 3**). MceB targets host mitochondria [38,39], but it also resides in the bacterial outer membrane [40]. Its retention in CEs that lack functional T4BSS machinery suggests that MceB may primarily function as a siderophore porin, although it may have acquired an additional role after *C. burnetii* diverged from CEs.

**Figure 3.**
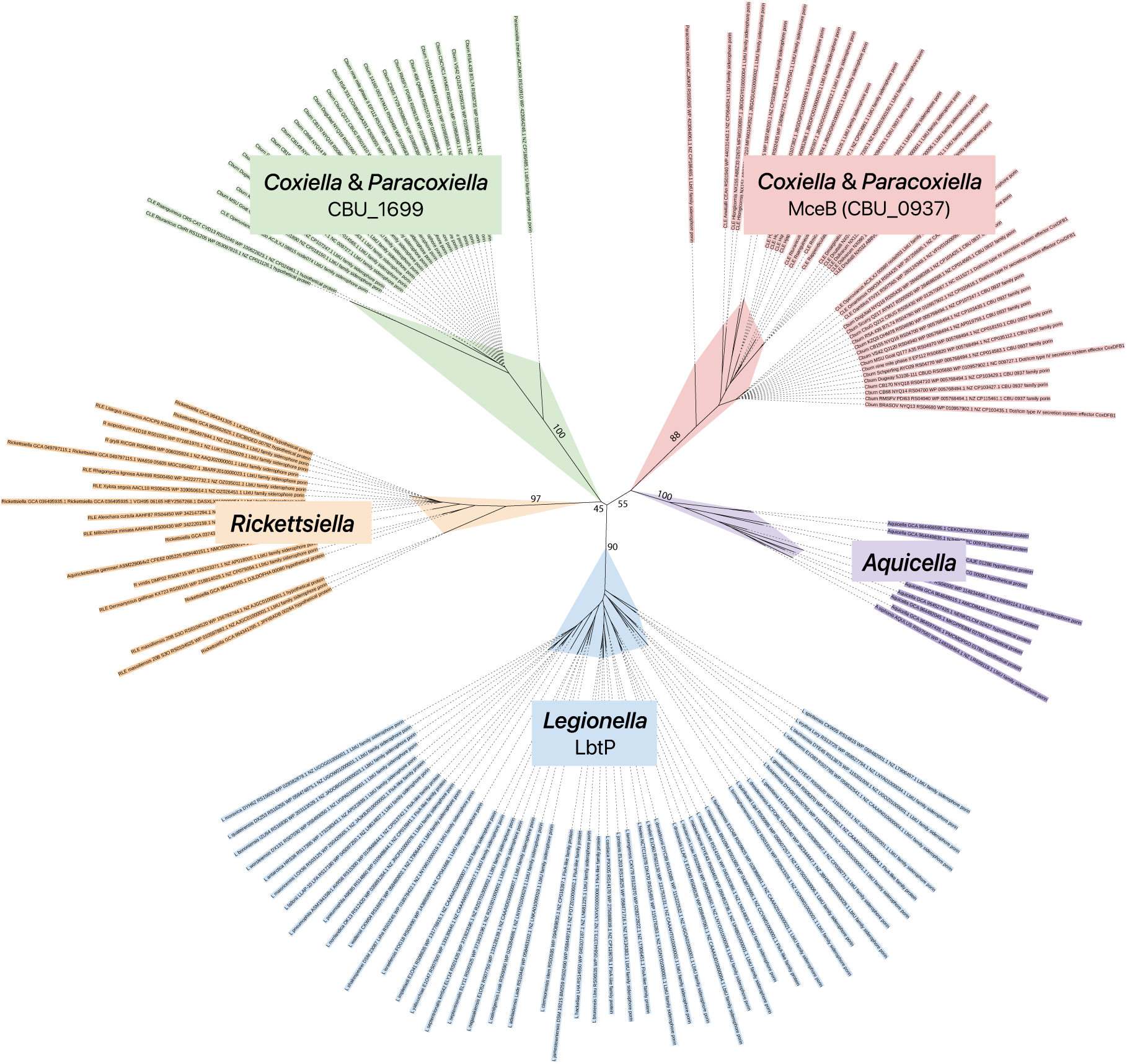
Protein phylogeny of LbtP-like sequences. The clade in red contains *C. burnetii* MceB and its orthologs found in *Paracoxiella* and all of the CEs. The clade in green contains *C. burnetii* CBU_1699 and its orthologs found in *Paracoxiella* and a few CEs. *Rickettsiella* and *Aquicella* each have only one LbtP-type gene. Bootstrap values are labeled for major branches.

CstK (CBU_0175), which affects endosomal trafficking and autophagy [41], may have been acquired through intra-Legionellales horizontal gene transfer (HGT). Outside *Coxiella* and *Paracoxiella*, the top-scoring matches to CstK were proteins from *Berkiella* species. Given the distant relationship between *Coxiella* and *Berkiella* (**Figure 1**), the high sequence identity (>65%), and the absence of CstK-like proteins in other Legionellales, these proteins are unlikely to be vertically inherited. Instead, the evidence suggests an HGT event either between *Coxiella* and *Berkiella* or from an unidentified common donor. The contrast between broadly conserved secretion machinery and more restricted effector distributions supports a model in which the T4BSS was inherited from an ancient Legionellales ancestor, while much of the *C. burnetii* effector repertoire has undergone lineage-specific expansion and innovation, likely reflecting adaptation to its distinct host range and intracellular niche.

### Recent assembly of O-antigen biosynthesis traits shaped a virulence-associated LPS structure

*C. burnetii* LPS contains the O-antigen-sugar virenose (**Figure 4**), whose loss prevents O-antigen elongation and results in attenuated virulence in mammals [42,43]. The pathway for virenose synthesis has not been fully characterized, but genes located in a genomic region lost from the avirulent NMII strain are known to participate in this process [23,25,27]. GDP-4-keto-6-deoxy-D-mannose, an intermediate in the biosynthesis of several O-antigen sugars such as fucose, perosamine, and rhamnose, is also thought to serve as an intermediate in virenose synthesis [44–46]. We found that the genes responsible for generating GDP-4-keto-6-deoxy-D-mannose — CBU_0294 (phosphomannomutase), CBU_0671 (GDP-mannose pyrophosphorylase), and CBU_0689 (GDP-mannose-4,6-dehydratase) — were likely inherited from the common ancestor of Coxiellaceae or an even earlier ancestor (**Figure 5**).

**Figure 4.**
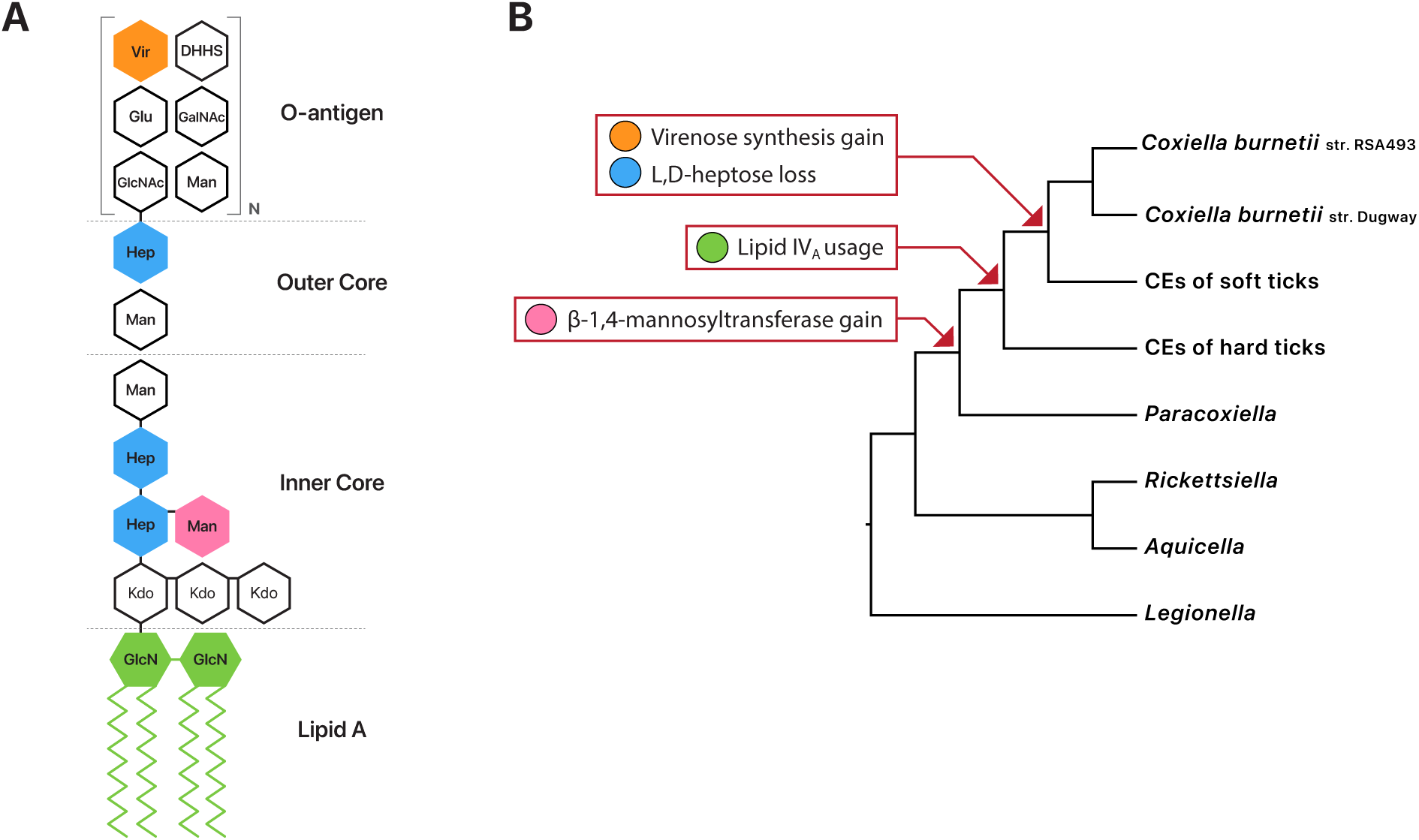
Summary of LPS structural changes in Coxiellaceae. (**A**) Components of LPS. (**B**) Gain and loss events.

**Figure 5.**
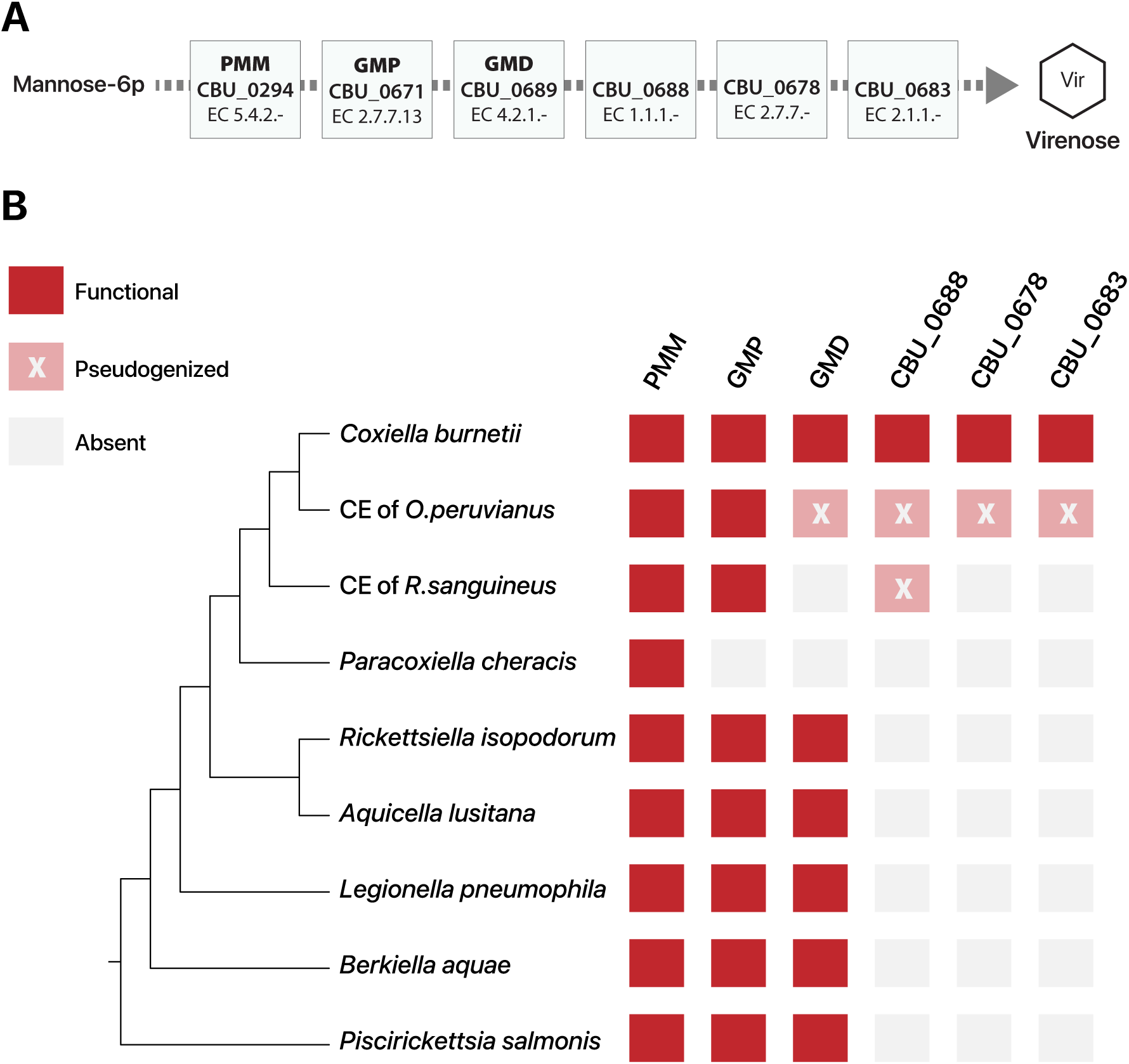
Distribution of genes likely required for virenose production. (**A**) Putative pathway for virenose synthesis. (**B**) Distribution of virenose synthesis genes across Legionellales. PMM: phosphomannomutase / phosphoglucomutase; GMP: GDP-mannose pyrophosphorylase; GMD: GDP-mannose-4,6-dehydratase.

GDP-4-keto-6-deoxy-D-mannose may function as the substrate for CBU_0688, a fucose synthase-like gene, and CBU_0683, a methyltransferase that has been proposed to participate in virenose synthesis [47,48]. Pseudogenized orthologs of CBU_0688 were detected in CEs of hard and soft ticks, whereas pseudogenized orthologs of CBU_0683 were identified only in *Ornithodoros* CEs, suggesting that at least part of the virenose synthesis pathway was present in the common ancestor of *C. burnetii* and CEs (**Figure 5**). Another gene experimentally shown to be essential for virenose synthesis is CBU_0678 [27]. Although its precise role remains unclear, CBU_0678 is similar to the bifunctional gene HldE involved in LPS inner-core sugar biosynthesis and our analysis indicates that it was likely acquired from Alphaproteobacteria (**Figure 6**). These distributions place several genes implicated in virenose production in the recent ancestry of *C. burnetii*, consistent with stepwise assembly of O-antigen biosynthetic capacity before the emergence of the fully virulent *C. burnetii* LPS structure.

**Figure 6.**
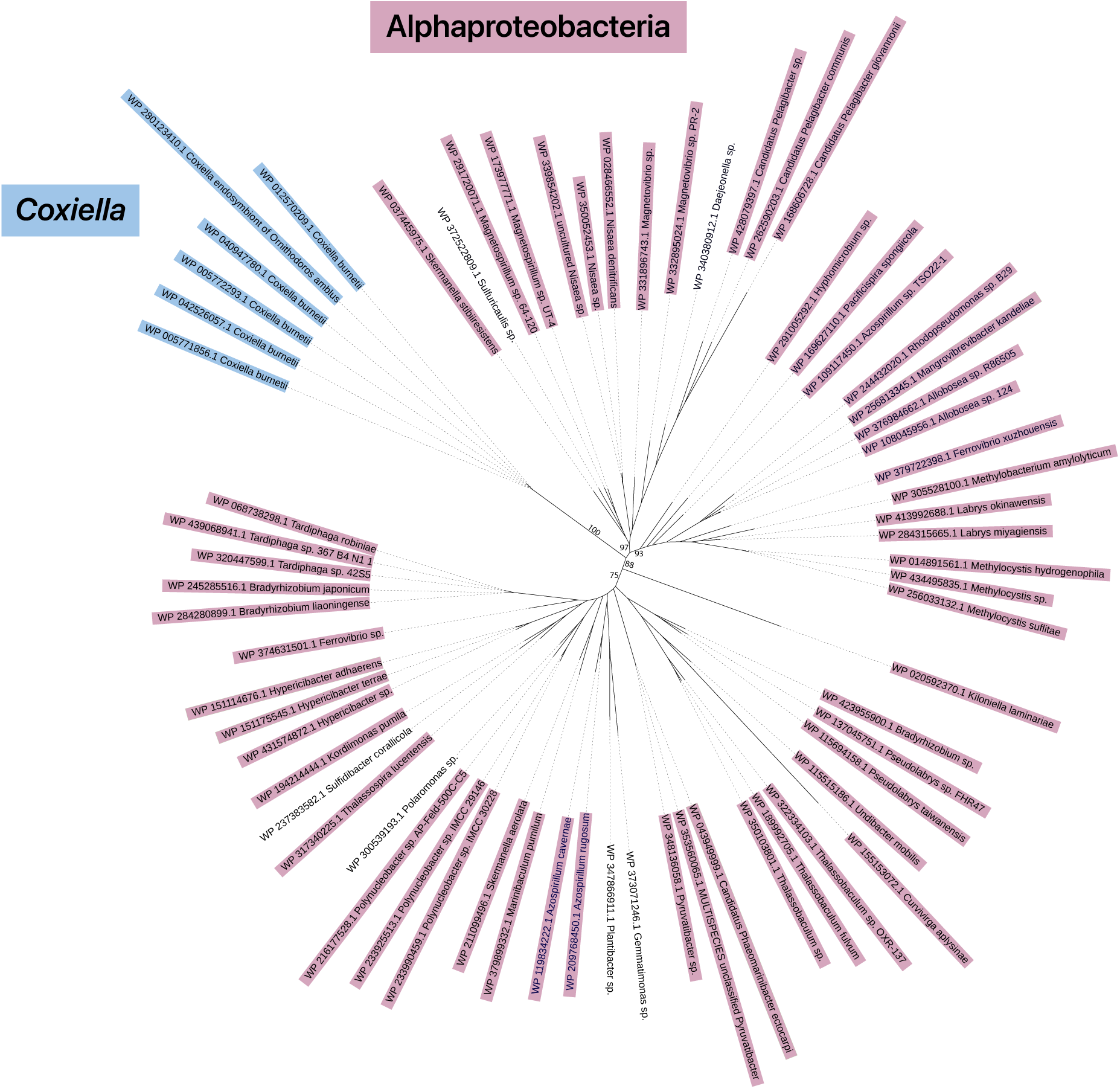
CBU_0678 was likely acquired from Alphaproteobacteria. Bootstrap values are labeled for major branches. Mauve: Alphaproteobacteria; Blue: *C. burnetii* and *Ornithodoros* CEs.

### LPS inner-core remodeling likely altered envelope properties and host recognition

The LPS core oligosaccharide links the lipid A anchor to the repeating O-antigen polysaccharide (**Figure 4**). It is generally less variable than the O-antigen and commonly contains the sugar L-glycero-beta-D-manno-heptose (LD-heptose). GmhD, which catalyzes the final step of the heptose synthesis pathway that converts DD-heptose to LD-heptose, has been lost in *C. burnetii* and *Ornithodoros* CEs (**Figure 7**). GmhD is retained in most hard-tick CEs and is also present in *Paracoxiella*, indicating that it was present in the common ancestor of *Coxiella*. Similar to *C. burnetii* LPS, a shift from LD-heptose to DD-heptose appears to have occurred independently in *Aquicella* and *Legionella* species (**Figure 7**). Changes in heptose stereochemistry could alter host recognition because bacterial heptose and heptose-derived metabolites are detected by innate immune pathways. The cytosolic ALPK1 pathway responds to bacterial ADP-heptose metabolites, and pulmonary surfactant protein D can bind heptose-containing LPS core structures; in both cases, recognition can differ between LD- and DD-heptose forms [49,50].

**Figure 7.**
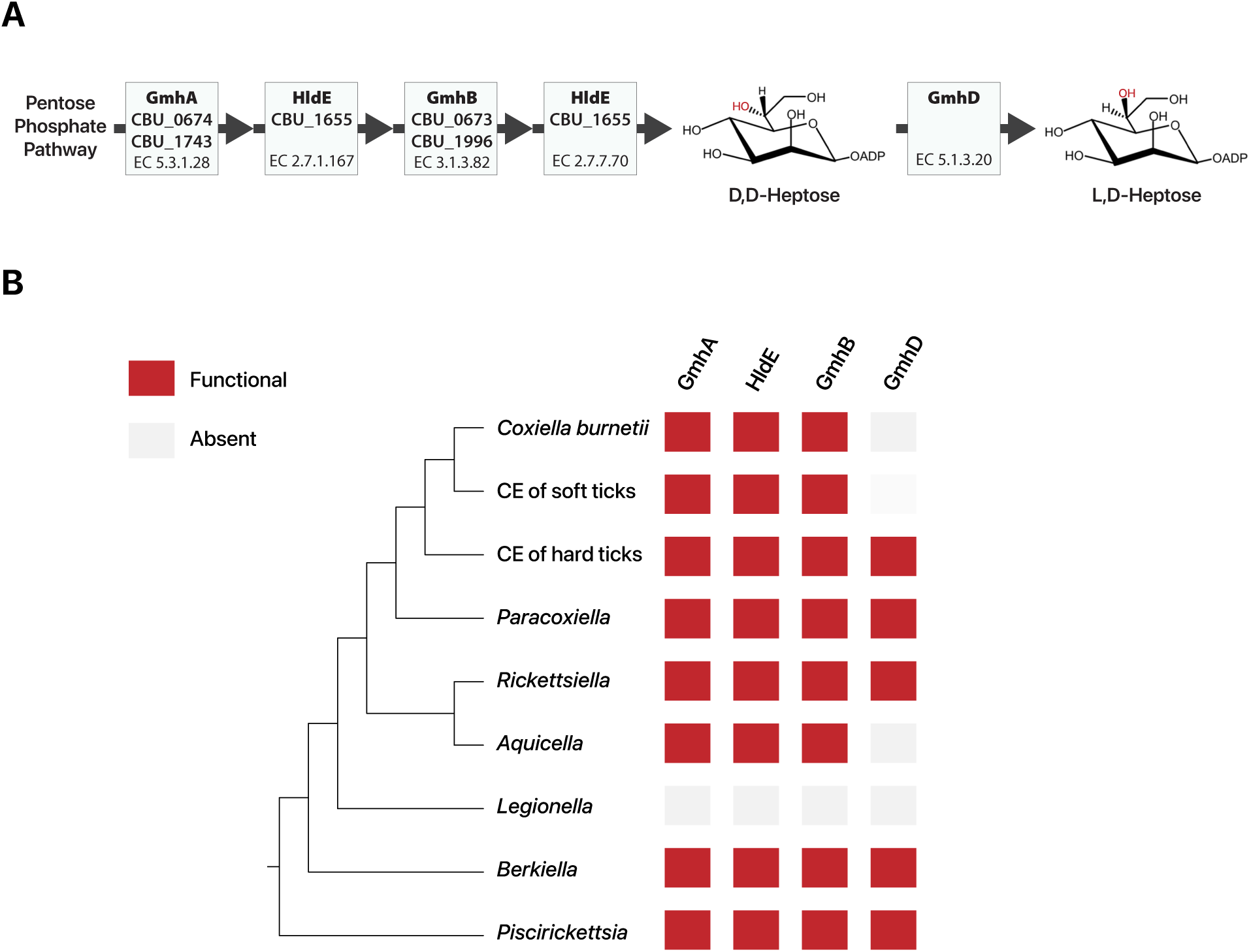
Synthesis of LD-heptose was lost in the common ancestor of *C. burnetii* and *Ornithodoros* CEs. (**A**) Pathway for generating L,D-Heptose. (**B**) GmhD, the enzyme responsible for converting DD-heptose to LD-heptose, is absent in *C. burnetii*.

*C. burnetii* LPS features a mannose substitution at the 4′-position of inner core heptose I (**Figure 4**), catalyzed by the glycosyltransferase CBU_1657 [27]. CBU_1657 was likely acquired in the common ancestor of *Coxiella* and *Paracoxiella* and has been retained in several hard- and soft-tick CEs. Phylogenetic analysis suggests that CBU_1657 was derived from Thermodesulfobacteriota, a phylum of extremophilic bacteria not known to include pathogens (**Figure 8**). *Geobacter sulfurreducens*, a member of this group, also contains mannose attached to HepI in its LPS inner core [51], suggesting that the mannosyltransferase activity of CBU_1657 originated prior to its acquisition via HGT by the common ancestor of *Coxiella* and *Paracoxiella*. Because HepI/HepII substitutions can influence serum sensitivity and interactions with host factors, acquisition of a mannosyltransferase provides a second mechanism by which inner-core remodeling could affect the bacterial surface [52]. The inferred loss of GmhD and acquisition of CBU_1657 support a model in which the *Coxiella* LPS inner core was reshaped through both gene loss and horizontal gene gain, potentially altering surface properties relevant to host interaction.

**Figure 8.**
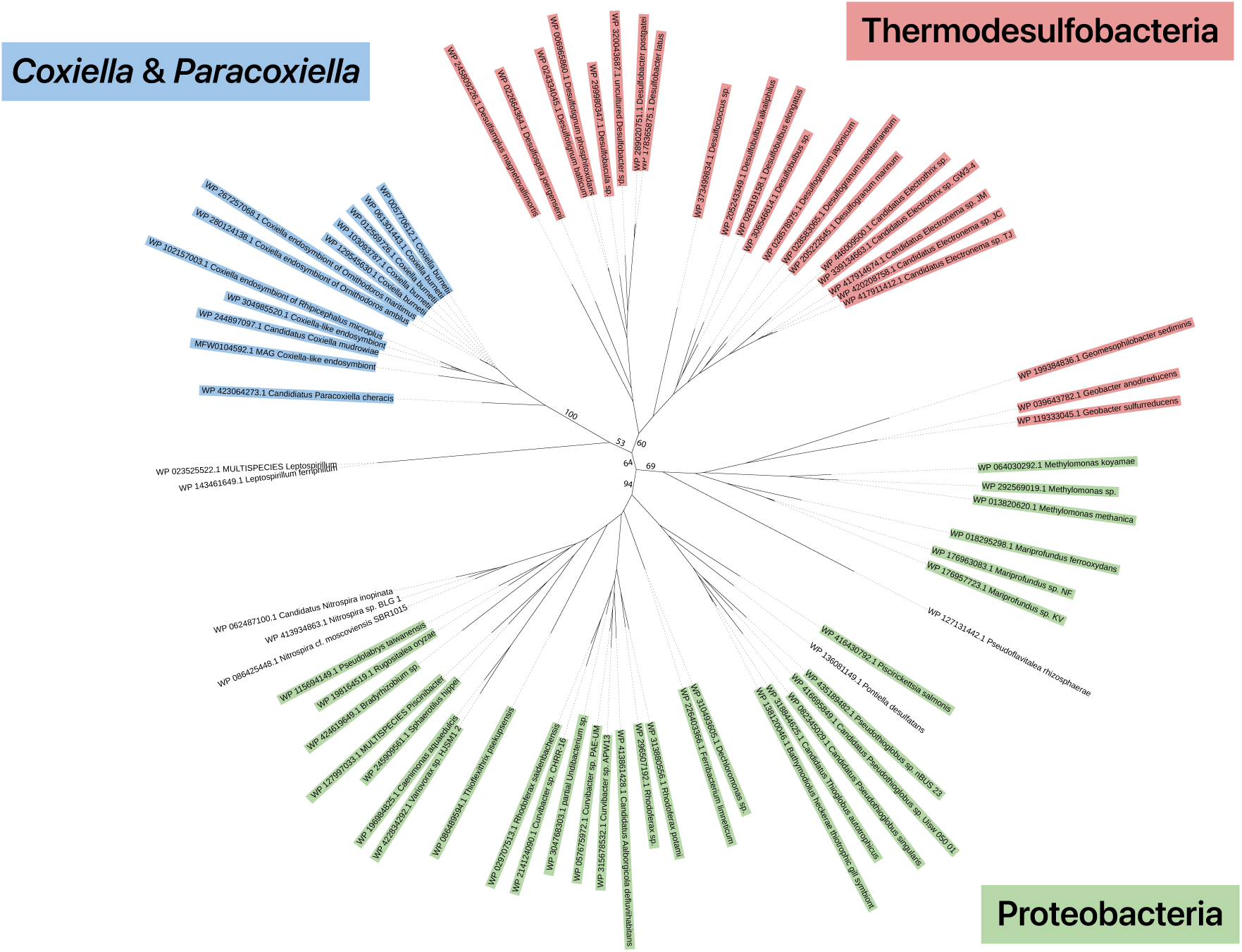
β-1,4 manosyltransferase (CBU_1657) may have been acquired from Thermodesulfobacteria. Bootstrap values are labeled for major branches. Blue: *Coxiella* and *Paracoxiella*; Red: Thermodesulfobacteria; Green: Proteobacteria.

### *C. burnetii* inherited a simplified lipid A anchor with weak TLR4-stimulatory potential

*C. burnetii* and the CEs appear to have the simplest form of lipid A, lipid IV_A_, which is tetra-acylated and requires only the core cytoplasmic lipid A biosynthesis enzymes (**Table 3**). In humans, lipid IV_A_ acts as a TLR4-MD2 antagonist and is only weakly agonistic in mice [53]. Unlike *Coxiella*, all other Legionellales encode one or more enzymes that carry out periplasmic modifications of lipid IV_A_ after its synthesis (**Table 3**). The distribution of these periplasmic modification genes suggests that the common ancestor of *Coxiella* already used a simplified lipid IV_A_ form as its LPS anchor, although this state may date to an even deeper ancestor. By retaining a simplified lipid IV_A_-like anchor, *C. burnetii* may have preserved an ancestral LPS feature that limits TLR4 stimulation, linking lipid A evolution to a functional property relevant to host interaction.

**Table 3.** Distribution of periplasmic lipid A modification genes across Legionellales.

|  | Cellular<br>Localization | *Coxiella*<br>*burnetii* | CEs | *Paracoxiella*<br>*cheracis* | *Rickettsiella*<br>*isopodorum* | *Aquicella*<br>*siphonis* | *Legionella*<br>*pneumophila* | *Berkiella*<br>*cookevillensis* |
| --- | --- | --- | --- | --- | --- | --- | --- | --- |
| **LpxA** | Cytosolic | 1 | 1 | 1 | 1 | 1 | 2 | 1 |
| **LpxC** | Cytosolic | 1 | 1 | 1 | 1 | 1 | 1 | 1 |
| **LpxD** | Cytosolic | 1 | 1 | 1 | 1 | 1 | 3 | 1 |
| **LpxH** | Cytosolic | 1 | 1 | 1 | 1 | 1 | 1 | 1 |
| **LpxB** | Cytosolic | 1 | 1 | 1 | 1 | 1 | 2 | 1 |
| **LpxK** | Cytosolic | 1 | 1 | 1 | 1 | 1 | 1 | 1 |
| **KdtA** | Cytosolic | 1 | 1 | 1 | 1 | 1 | 1 | 1 |
| **lpxL/P** | Periplasmic | 0 | 0 | 0 | 1 | 0 | 3 | 0 |
| **LpxO** | Periplasmic | 0 | 0 | 0 | 1 | 0 | 0 | 0 |
| **ArnT<sup>a</sup>** | Periplasmic | 0 | 0 | 0 | 1 | 1 | 0 | 0 |
| **PagP** | Periplasmic | 0 | 0 | 0 | 0 | 0 | 1 | 0 |
| **PagL** | Periplasmic | 0 | 0 | 2 | 1 | 1 | 0 | 1 |

<sup>a</sup> in addition to other *arn* genes

### Horizontal acquisition of polyamine biosynthesis expanded stress-tolerance potential

Polyamines such as putrescine and homospermidine contribute to bacterial tolerance of high temperature, reactive oxygen species, and low pH [54–56]. *C. burnetii*’s polyamine synthesis gene cluster (CBU_0720-CBU_0722) is also present in CEs, *Paracoxiella*, and *Rickettsiella*, but not in other Legionellales. This distribution suggests that the cluster was acquired in the common ancestor of Coxiellaceae, and our phylogenetic analysis indicates that it was horizontally acquired from Alphaproteobacteria (**Figure 9**). Experiments in alphaproteobacterial family Pelagibacteraceae indicate that although the type III PLP-dependent decarboxylase encoded in this cluster (CBU_0722) is structurally similar to an ornithine decarboxylase, it functions predominantly as an arginine decarboxylase [57] (**Figure 10**). Assuming this is also true in *Coxiella*, this gene cluster constitutes a complete pathway for the conversion of arginine to putrescine and its derivative homospermidine. These data point to horizontal acquisition of a polyamine biosynthesis pathway whose predicted products could improve *C. burnetii*’s tolerance to oxidative or acidic stress in its intracellular niche.

**Figure 9.**
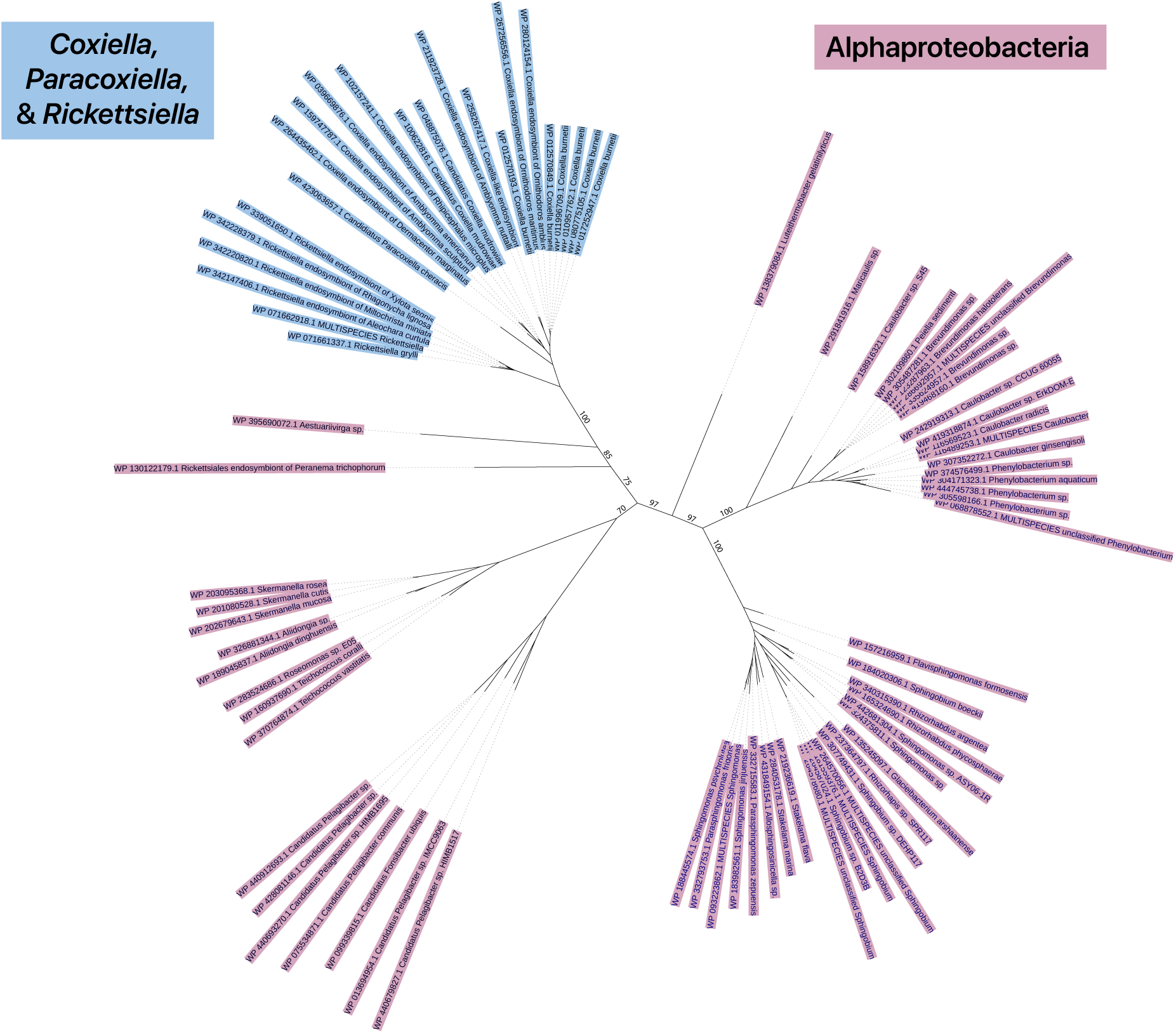
Coxiellaceae appears to have acquired a polyamine synthesis gene cluster (CBU_0720-22) from Alphaproteobacteria. *Coxiella*, *Paracoxiella*, and *Rickettsiella* are highlighted in blue while Alphaproteobacteria are highlighted in mauve. A representative ML tree of CBU_0722 is shown. ML trees of CBU_0720 and CBU_0721 produced similar results. Bootstrap values are labeled for major branches.

**Figure 10.**
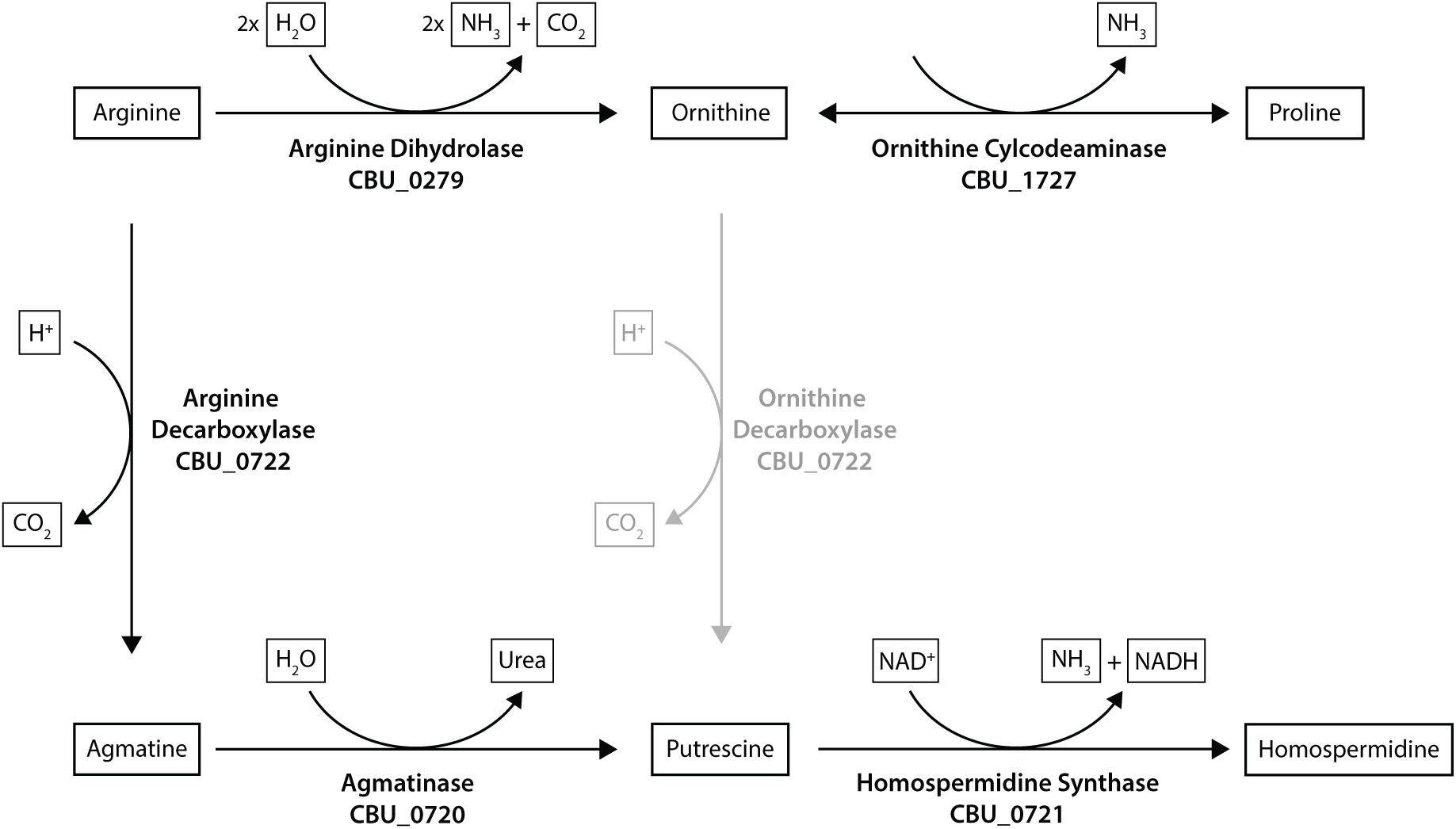
Pathways for polyamine synthesis and arginine metabolism in *C. burnetii*.

### Acquired ammonia-producing enzymes connect arginine metabolism to pH homeostasis

*C. burnetii*’s pathway for converting arginine to proline consists of two ammonia-producing enzymes (**Figure 10**). These atypical genes can increase pH in an acidic environment [58–60]. The first enzyme (CBU_0279) is homologous to the N-terminal domain of arginine dihydrolase, which is characteristically found in cyanobacteria [61] (**Figure 11**). Three *Legionella* taxa also encode genes homologous to arginine dihydrolase, but the sparse distribution of this function across Legionellales suggests that these genes are unlikely to be orthologous. The second enzyme, ornithine cyclodeaminase (CBU_1727), was also likely acquired in the common ancestor of *Coxiella* and *Paracoxiella*. BLASTp analysis identified homologs in several distantly related proteobacteria and a CBU_1727 protein tree placed its closest relative in the archaeal genus *Thermofilum* (**Figure 12**). By adding atypical ammonia-producing steps to arginine metabolism, the *C. burnetii* lineage may have gained a mechanism that could contribute to pH homeostasis in acidic environments.

**Figure 11.**
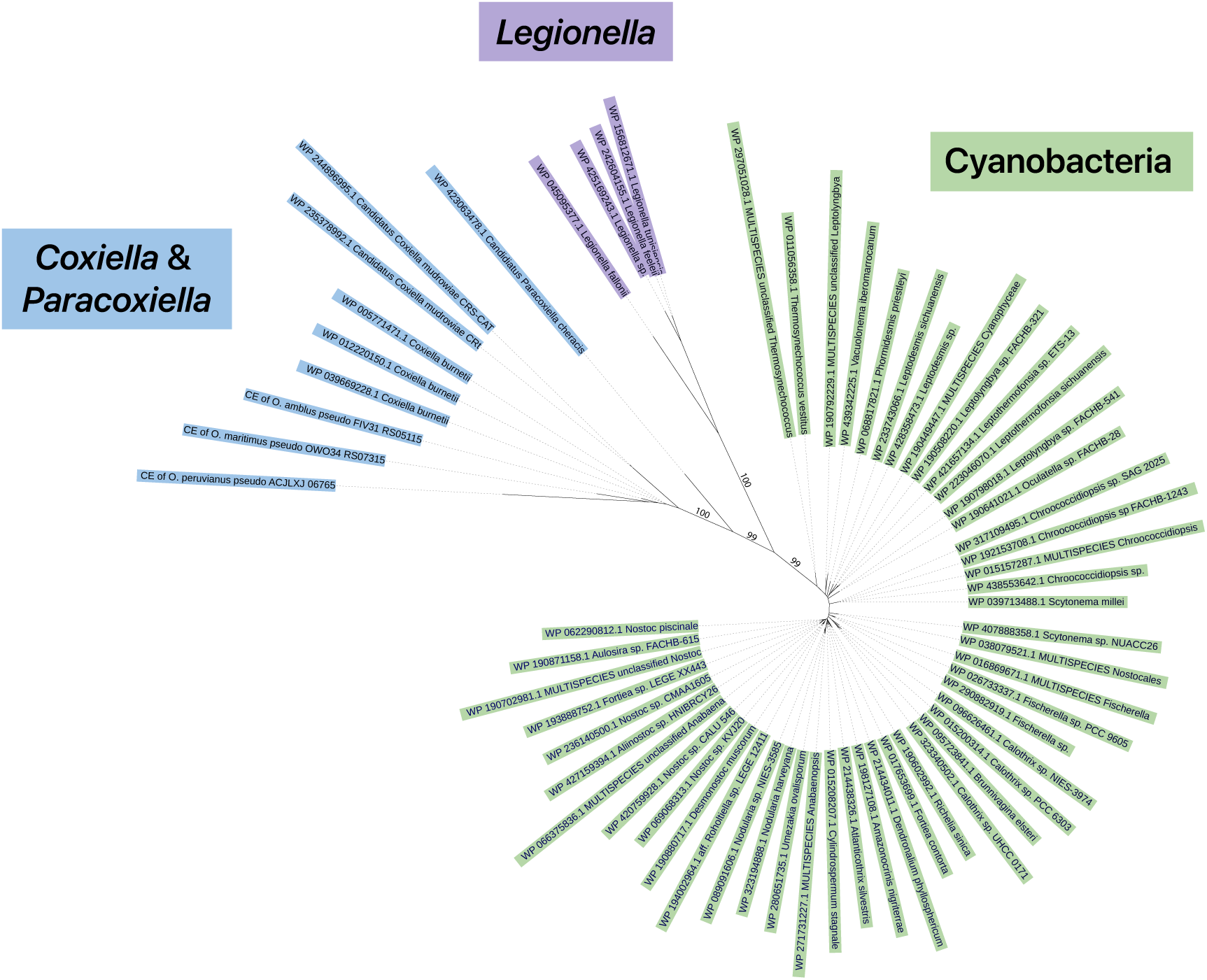
Arginine dihydrolase (CBU_0279) was likely acquired in the common ancestor of *Coxiella* and *Paracoxiella* (blue) from cyanobacteria (green). Homologs were also found in three *Legionella* species (purple). Bootstrap values are labeled for major branches.

**Figure 12.**
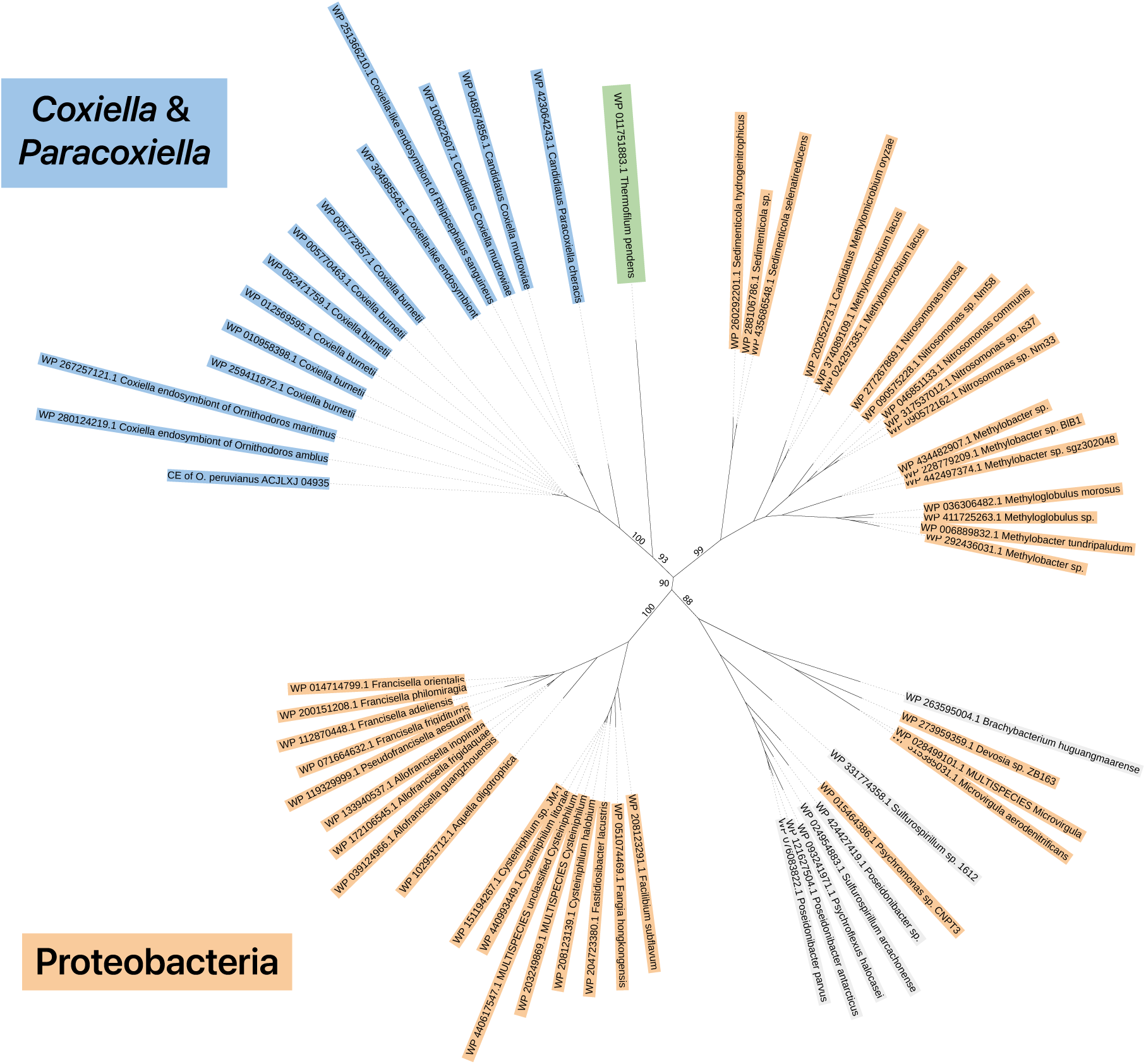
Ornithine cyclodeaminase (CBU_1727) may have been horizontally acquired. *Coxiella* and *Paracoxiella* are shown in blue. The closest homolog was found in an archaeon (green), while most others were in various proteobacteria (orange). Bootstrap values are labeled for major branches.

### Fatty acid pathway remodeling expanded membrane stress-adaptation capacity

Since diverging from *Legionella*, the lineage leading to *C. burnetii* has undergone several changes in fatty acid synthesis and modification that likely affect stress tolerance [62] (**Figure 13**). *Coxiella* is the only Legionellales lineage predicted to produce monounsaturated fatty acids (UFAs) using FabA (CBU_0037) and FabB (CBU_0035). FabAB is part of a larger lipid metabolism gene cluster whose distribution suggests that it was acquired in the common ancestor of *Coxiella*, as these genes are found only in *C. burnetii* and the CEs (**Figure 14**). Most of these genes have older functional counterparts elsewhere in the genome; thus, *C. burnetii* encodes two fatty acid synthesis systems: an ancestral pathway for saturated fatty acids and a more recently acquired pathway that can generate UFAs. Low pH typically increases the density and order of lipid bilayers, so genes that can maintain membrane fluidity could be beneficial in this environment [63]. *C. burnetii*’s membrane is primarily composed of branched-chain fatty acids [64,65], so the specific role of UFAs is unclear, but likely allows *C. burnetii* to fine tune membrane fluidity in response to various environmental stressors including pH.

**Figure 13.**
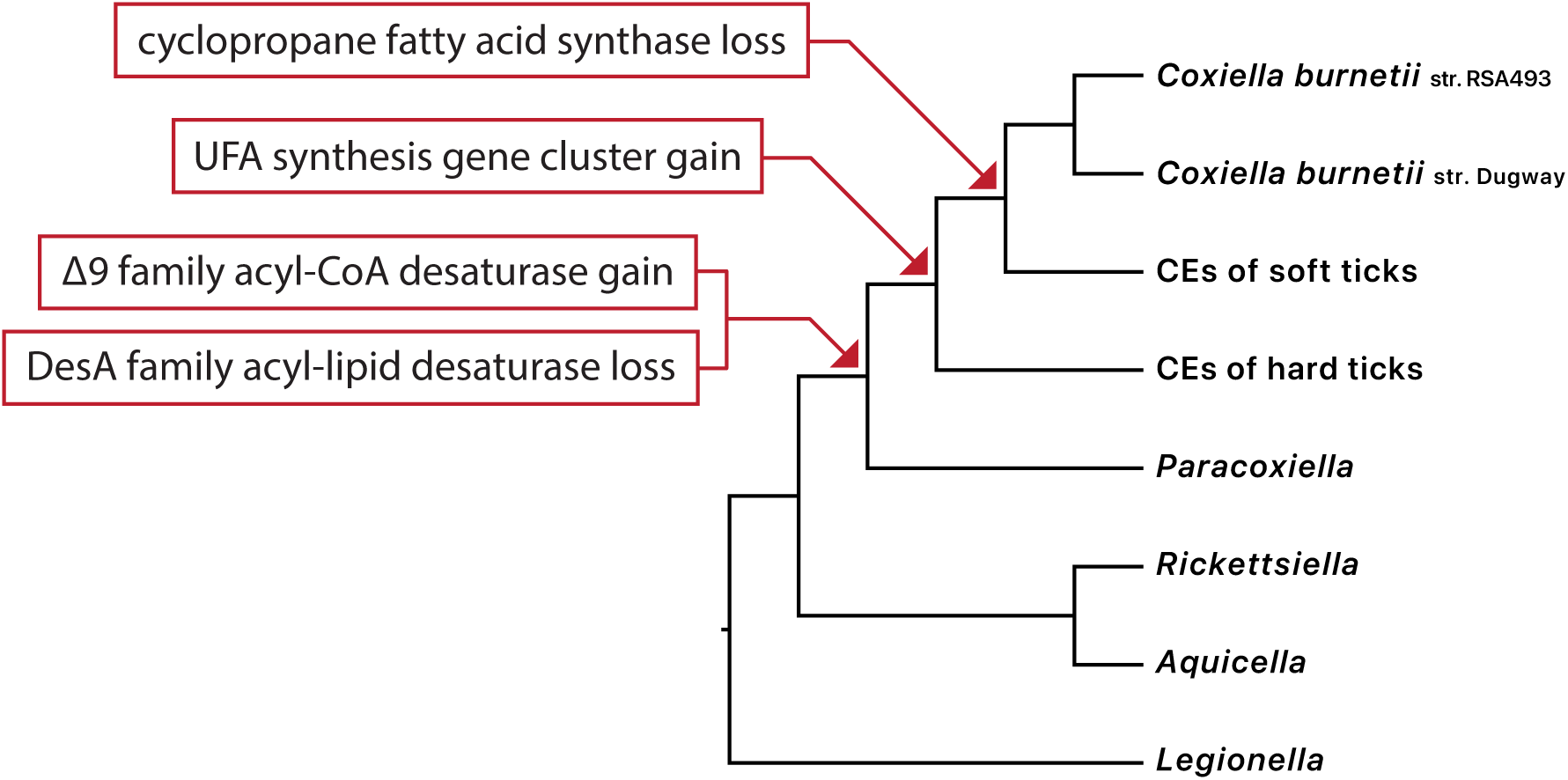
Summary of changes to fatty acid synthesis and modification in the lineage leading to *C. burnetii*.

**Figure 14.**
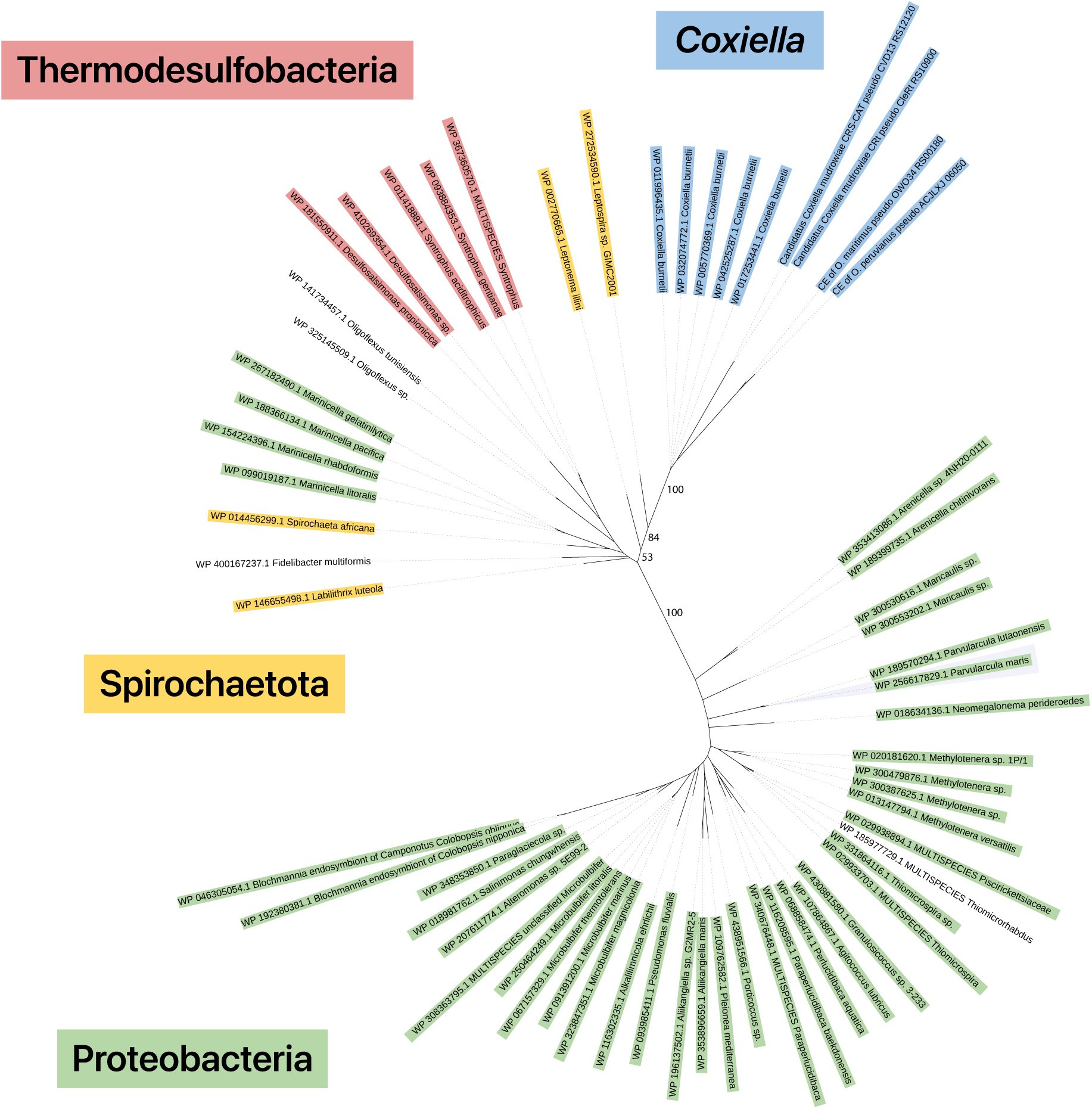
Potential horizontal acquisition of FabA (CBU_0037) in the common ancestor of *Coxiella*. Closest homologs of *Coxiella’*s FabA (blue) are found in a diverse group of bacteria including Proteobacteria (green), spirochetes (yellow), and Thermodesulfobacteriota (red). Bootstrap values are labeled for major branches.

The common ancestor of *Coxiella* and *Paracoxiella* also gained a delta-9 acyl-CoA desaturase (CBU_0920), which desaturates fatty acyl chains bound to acyl-CoA. That same ancestor lost a DesA-family acyl-lipid desaturase that acts on fatty acyl chains already incorporated into membrane lipids. Unlike many other Legionellales, *C. burnetii* also lacks cyclopropane fatty acid synthase (CfaS), and our reconstruction suggests that this gene was lost in the common ancestor of *C. burnetii* and *Ornithodoros* endosymbionts. Collectively, this pattern of gene loss and gain suggests that fatty acid desaturation became more closely coupled to fatty acid synthesis during *Coxiella* evolution, potentially expanding the capacity to regulate membrane composition under environmental stress.

### Recent acquisition of the Mrp antiporter may support pH homeostasis

*C. burnetii* encodes a multi-subunit Mrp (Multiple resistance and pH adaptation) antiporter gene cluster that appears to be essential because multiple attempts to generate mutants have been unsuccessful [66,67]. Additionally, Mrp genes have been pseudogenized or lost in CEs, indicating that they are not critical in tick symbionts that do not inhabit acidic intracellular vacuoles. The Mrp cluster was likely acquired in the common ancestor of *C. burnetii* and the *Ornithodoros* CEs (**Figure 15**). Mrp antiporters are also present in a small number of *Legionella* species, but their distribution across Legionellales suggests that the system was acquired independently in *Coxiella* and *Legionella*. The function of *C. burnetii’*s Mrp system has not been experimentally established but is thought to aid in pH homeostasis because it is required for growth in both acidic axenic medium and within host cells [66–68]. Mrp is typically associated with alkaliphilic and halophilic bacteria, where protons are imported in exchange for Na^+^/Li^+^ export [69,70]. However, Mrp mutation in *Pyrococcus furiosus* was found to cause growth impairment under acidic conditions [71], suggesting that the antiporter likely contributes to pH homeostasis during growth within the acidic CCV.

**Figure 15.**
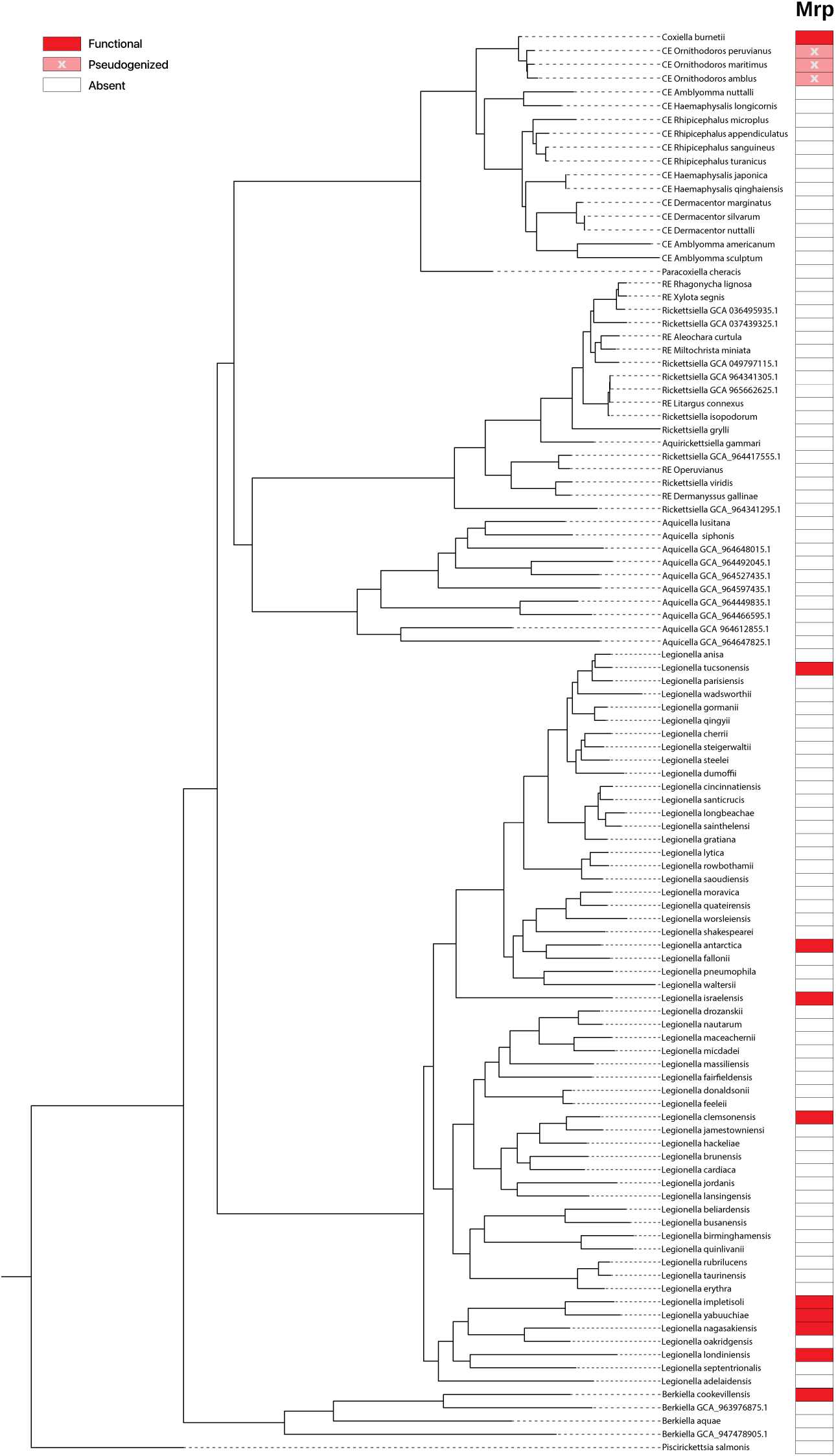
Distribution of Mrp cation/proton transporter genes across Legionellales.

### An acquired mannose catabolism pathway connects host-derived carbon to central metabolism

Our analyses indicate that the common ancestor of Coxiellaceae gained a gene cluster (CBU_1275-CBU_1277) that may allow host-derived mannose to be used as a carbon and energy source (**Figure 16**). This pathway resembles hexuronate catabolism pathways found in bacteria and fungi, but it uses an aldohexose dehydrogenase (AldT, CBU_1276) to generate mannonate directly from mannose rather than from a hexuronate. A mannose utilization pathway of this general type has been characterized in the archaeon *Thermoplasma* [72]. A conserved domain search further indicates that CBU_1276 is similar to a gluconate 5-dehydrogenase-like short-chain dehydrogenase/reductase (E-value 1e-47), which is consistent with the proposed function. Mannose catabolism is widespread in bacteria; however, it generally involves converting mannose to the glycolytic intermediate fructose-6-phosphate. The unusual method in Coxiellaceae instead converts mannose to pyruvate and to our knowledge has not been characterized outside of archaea. Although the physiological role of this pathway remains to be experimentally validated, it may enable *C. burnetii* to utilize host-derived mannose through a distinct metabolic route that feeds directly into central carbon metabolism.

**Figure 16.**
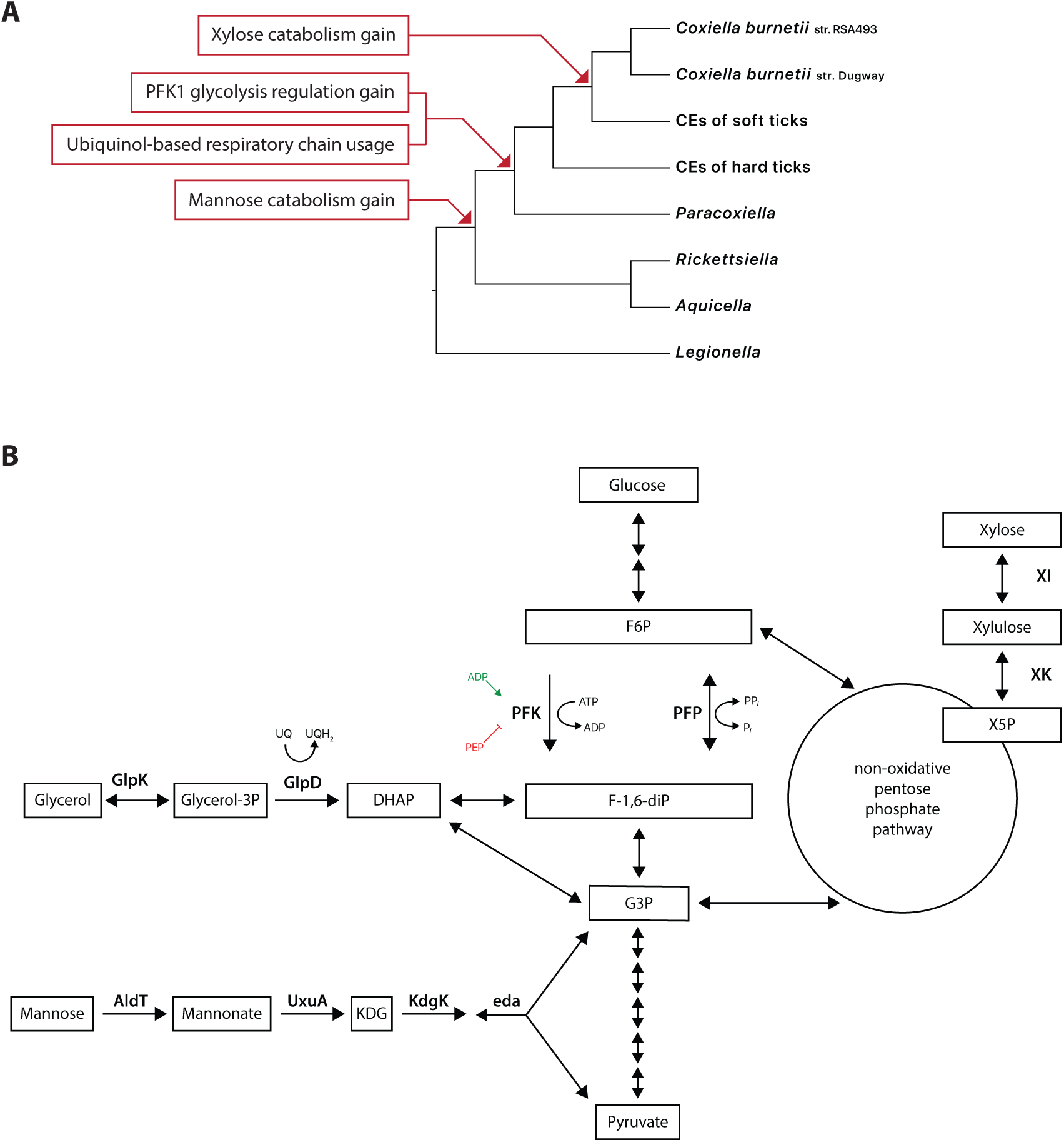
Changes in sugar catabolism, glycolytic control, and respiratory metabolism. (**A**) Summary of gene gain events that have affected sugar utilization in the lineage leading to *C. burnetii*. (**B**) Sugar and sugar alcohol catabolism pathways in *C. burnetii*. AldT: aldohexose dehydrogenase; DHAP: dihydroxyacetone phosphate; eda: 2-dehydro-3-deoxyphosphogluconate aldolase; F-1,6-diP: Fructose-1,6-bisphosphate; F6P: Fructose-6-phosphate; G3P: glyceraldehyde 3-phosphate; KDG: 2-keto-3-deoxygluconate; KdgK: 2-dehydro-3-deoxygluconokinase; PFK: ATP-dependent phosphofructokinase; PFP: inorganic pyrophosphate-phosphofructokinase; UQ: ubiquinone; UQH2: ubiquinol; UxuA: mannonate dehydratase; X5P: xylulose 5-phosphate; XI: xylose isomerase; XK: xylulokinase.

### Xylose utilization was assembled from older transport and newer catabolic functions

*C. burnetii* encodes a xylose isomerase (CBU_0344a) and a xylulose kinase (CBU_0346), which likely convert imported xylose to xylulose-5-phosphate, a pentose phosphate pathway intermediate (**Figure 16**). Pseudogenized versions of these genes were identified in *Ornithodoros* CEs, whereas no orthologs were detected in other Legionellales, indicating that this cluster was likely acquired in the common ancestor of *C. burnetii* and *Ornithodoros* CEs. The adjacent sugar transporter (CBU_0347), annotated as a xylose-proton symporter, appears to be older and likely dates to the common ancestor of Coxiellaceae and *Legionella*. The evolutionary history of this locus suggests that xylose utilization was assembled in stages, with recent acquisition of catabolic enzymes building on an older sugar transport system and potentially creating a route for host-derived sugars to enter the pentose phosphate pathway in *C. burnetii*.

### Acquired ATP-dependent phosphofructokinase added regulatory control to glycolysis

In addition to the inorganic pyrophosphate phosphofructokinase (PFP, CBU_1273) found throughout the Legionellales, the common ancestor of *Coxiella* gained an ATP-dependent phosphofructokinase (PFK, CBU_0341). The ancestral bidirectional PFP can function in both glycolysis and gluconeogenesis, whereas the newly-acquired PFK reaction is generally considered irreversible and therefore only glycolytic (**Figure 16**). PFKs also typically contain an allosteric site, which allows tighter regulation of pathway flux [73], thereby likely enhancing *C. burnetii*’s regulatory control over glycolysis. Acquisition of ATP-dependent PFK added a second phosphofructokinase with biochemical properties distinct from the ancestral PFP, potentially allowing *C. burnetii* to direct carbon flux toward glycolysis with greater regulatory control as nutrient availability fluctuates within the CCV.

### Respiratory-chain simplification reduced reliance on ROS-prone cytochrome c systems

The common ancestor of *Coxiella* and *Paracoxiella* lost all cytochrome c-associated systems, including the cytochrome caa_3_ terminal oxidase, cytochrome bc_1_ complex, and cytochrome c maturation machinery that are present in most other Legionellales (**Figure 17**). These systems can be bypassed because *Coxiella* and *Paracoxiella* encode ubiquinol terminal oxidases that accept electrons directly from NADH dehydrogenase and succinate dehydrogenase. *Coxiella*, *Paracoxiella*, and *Aquicella* encode a cytochrome bo_3_ ubiquinol oxidase (CBU_1035d, CBU_1038-CBU_1040), suggesting that this system was acquired in the common ancestor of Coxiellaceae. Unlike the ancestral cytochrome bd ubiquinol oxidase, cytochrome bo_3_ acts as a proton pump [74]. Cytochrome bo_3_ is the only terminal oxidase retained in all CEs, possibly because proton pumping remains advantageous even in the symbiotic lifestyle. Acquisition of cytochrome bo_3_ may have reduced the selective pressure to maintain cytochrome c-dependent systems. Loss of those systems may in turn benefit *C. burnetii* in its present environment because the cytochrome bc_1_ complex is prone to generating reactive oxygen species when challenged by a steep proton gradient [75,76]. Loss of cytochrome c-dependent systems alongside retention of ubiquinol terminal oxidases indicates a shift toward a streamlined respiratory chain that may preserve energy conservation while reducing dependence on complexes that could be costly in the acidic CCV environment.

**Figure 17.**
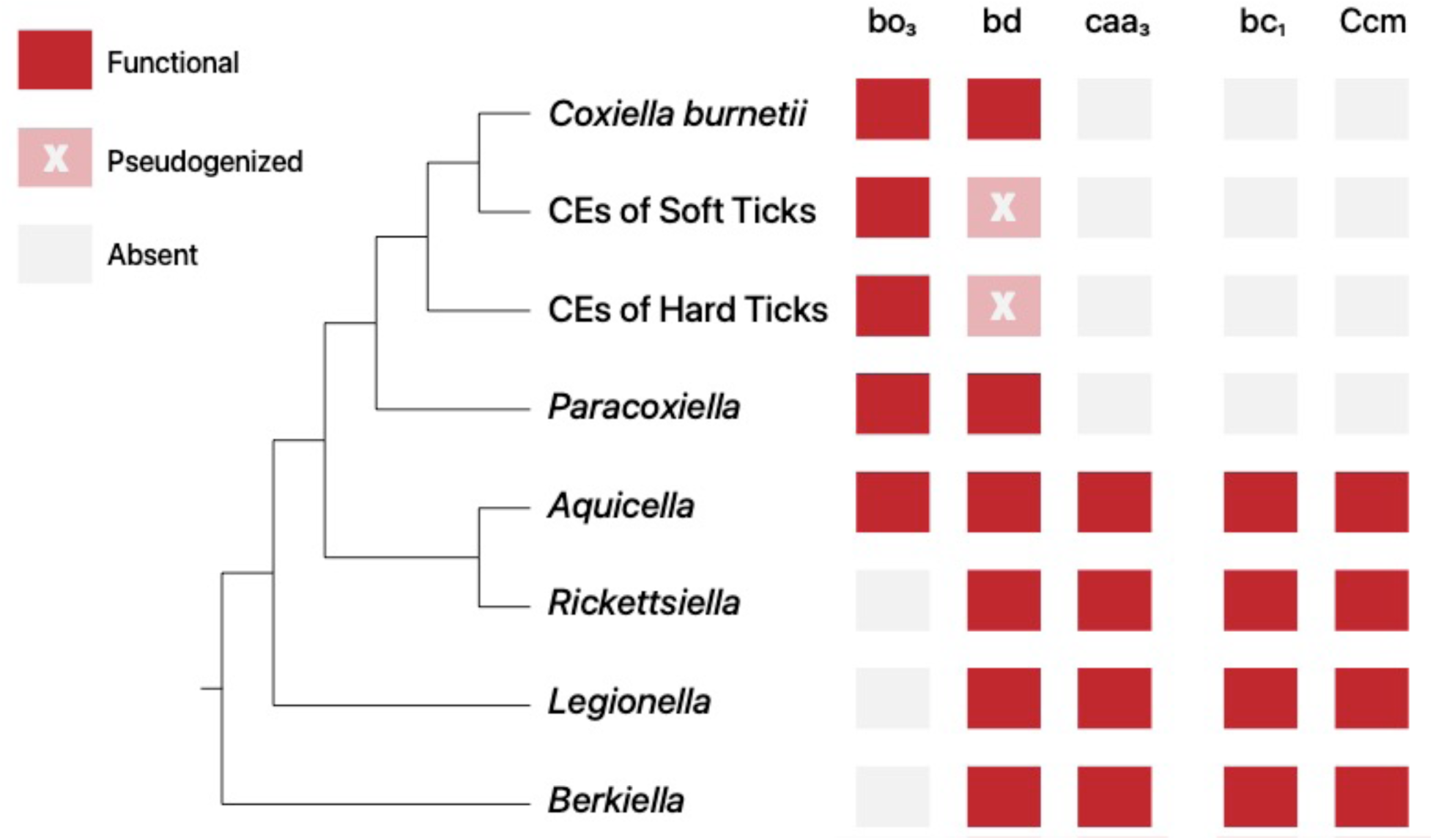
Distribution of cytochrome genes in Legionellales. bo_3_: cytochrome bo_3_ ubiquinol oxidase; bd: cytochrome bd ubiquinol oxidase; caa_3_: cytochrome caa_3_ oxidase; bc_1_: cytochrome bc_1_ complex; Ccm: cytochrome c maturation machinery.

## DISCUSSION

Our analyses support a model in which vertebrate pathogenicity in *C. burnetii* emerged through the stepwise assembly and remodeling of traits within an ancestral pathogenic background. Earlier phylogenomic work showed that soft-tick *Coxiella* endosymbionts retain remnants of a more virulent ancestor, placing *C. burnetii* within a broader evolutionary history of host association [32]. Building on that framework, our trait-by-trait reconstruction of gene gain and loss shows that key infection-associated traits trace to ancestors at different evolutionary depths. This Legionellales-wide analysis indicates that many systems central to *C. burnetii* pathogenesis, including traits associated with vertebrate infection and replication in the CCV, predate the modern pathogen and accumulated across multiple ancestral lineages.

Comparative studies of other bacterial pathogens have shown that virulence evolution commonly involves gene acquisition, gene loss, pseudogenization, genome rearrangement, and modification of preexisting cellular functions. In *Yersinia*, genomic studies have emphasized how mammalian pathogenesis evolved through a mixture of gene gain, gene loss, and genome rearrangement [77]. In *Shigella*, genomic plasticity and acquisition of a distinctive virulence plasmid enabled the repeated emergence of invasive pathogens from different *Escherichia coli* ancestors, while accelerated gene loss and pseudogenization accompanied specialization to a restricted host-associated niche [78,79]. *Salmonella* provides another example in which horizontally acquired pathogenicity islands encode secretion systems and effectors that are central to host interaction and invasion [80]. Work on *Vibrio cholerae* adds an ecological dimension, showing that environmental populations can harbor virulence-associated alleles, mobile elements, and pathogenic traits before the emergence of successful pathogenic clones [81–83].

These examples largely involve environmental, enteric, or facultatively host-associated bacteria, where access to diverse microbial communities can provide repeated opportunities for gene exchange. In obligately intracellular lineages, opportunities for horizontal gene transfer may be more episodic, depending on rare ecological contexts such as coinfection of the same host cell [84–86]. *C. burnetii* therefore provides an opportunity to examine how virulence-related traits accumulate in a specialized intracellular lineage. Our study shows that the modern *C. burnetii* virulence profile was shaped by familiar evolutionary processes, including HGT, gene loss, retention, duplication, and repurposing, but that these changes were layered across longer evolutionary timescales.

The evolutionary history of the Dot/Icm type IVB secretion system is consistent with this model. Most structural components of the secretion apparatus were already widespread across Legionellales, indicating that the machinery itself is ancient within the order. In contrast, the effector repertoire associated with *C. burnetii* appears to be much younger and concentrated within *Coxiella* and its closest relatives. The common ancestor shared with *Ornithodoros* endosymbionts likely encoded most *C. burnetii* effectors, implying that much of the host-manipulation capacity now associated with vertebrate infection was already present before the emergence of the modern pathogen. At the same time, the limited distribution of many effectors outside *Coxiella* and *Paracoxiella* suggests that effector repertoires remain evolutionarily labile and are shaped strongly by lineage-specific host associations. This pattern fits the broader view that secretion systems can be conserved over long periods whereas their cargo proteins turn over more rapidly as intracellular bacteria adapt to different eukaryotic environments.

Our reconstruction of lipopolysaccharide evolution similarly suggests that key immune-evasion traits were assembled in stages rather than acquired all at once. Several changes affecting the *C. burnetii* envelope appear to have accumulated along successive *Coxiella* ancestors, including lipid A simplification, modifications to the inner core, and acquisition of genes associated with synthesis of unusual O-antigen sugars. Inner-core remodeling provides one plausible route for altered host recognition. Loss of GmhD has shifted heptose stereochemistry from LD- to DD-heptose, which may reduce recognition by host factors that respond differently to heptose isomers [25,87]. In parallel, acquisition of CBU_1657 introduced a mannose substitution at HepI, and substitutions at HepI or HepII are broadly conserved among pathogenic bacteria and can influence serum susceptibility. The specific contribution of the *C. burnetii* mannose substitution remains unresolved, but these observations suggest that inner-core remodeling could affect both immune recognition and envelope interactions. The O-antigen shows a similar pattern of staged evolution: the common ancestor of *C. burnetii* and *Ornithodoros* endosymbionts may already have possessed several genes implicated in virenose synthesis, a distinctive sugar that is critical for full O-antigen elaboration and vertebrate virulence in *C. burnetii*. More generally, the inferred sequence of LPS changes argues against a single decisive innovation and instead supports progressive remodeling of the cell surface, with each step potentially altering resistance to host recognition, antimicrobial factors, or other environmental pressures.

Several traits associated with stress tolerance also appear to have accumulated before the emergence of modern *C. burnetii*. Horizontal acquisition of a polyamine biosynthesis pathway, addition of atypical ammonia-producing steps in arginine metabolism, and changes in fatty acid synthesis and desaturation all have plausible connections to survival under acidic, oxidative, or otherwise stressful conditions. These functions are not specific to the CCV, and some may originally have supported tolerance of broader environmental or host-associated stresses. This is important because the CCV, in addition to being acidic, is rich in lysosomal enzymes, reactive oxygen species, osmotic stress, and host-derived nutrients. The inferred history therefore suggests that *C. burnetii* acid tolerance may have emerged from a combination of general stress-resilience traits and more niche-relevant systems. In this context, the Mrp cation/proton antiporter is notable because it stands out as a more direct candidate for improved pH homeostasis. Its apparent essentiality in *C. burnetii*, coupled with loss or pseudogenization in tick endosymbionts, is consistent with the idea that it became especially important in the acidic intracellular compartment exploited by the pathogen. Together, these patterns suggest that growth in the acidic CCV was enabled by a layered set of traits, combining ancestral general stress-resilience functions with recently acquired systems more directly tied to pH homeostasis.

Carbon metabolism and respiration provide a parallel example of sequential trait assembly. The mature CCV is nutrient rich, and *C. burnetii* is able to use both amino acids and sugars as carbon and energy sources. Our analyses suggest that this flexibility reflects several distinct evolutionary events rather than one broad metabolic expansion. Coxiellaceae acquired an unusual mannose catabolism pathway predicted to route mannose-derived carbon into central metabolism. Xylose utilization appears to have been assembled in stages, with recent acquisition of catabolic enzymes building on an older sugar transporter that may have broader substrate specificity, including glucose transport [30]. The addition of ATP-dependent phosphofructokinase alongside the ancestral pyrophosphate-dependent enzyme may also have increased regulatory control over carbon flux through glycolysis. These changes occurred against a background of respiratory chain remodeling, including loss of cytochrome c-associated systems and reliance on ubiquinol terminal oxidases. This respiratory architecture eliminated dependence on cytochrome c-associated complexes that can generate reactive oxygen species under steep proton gradients but also reduces the overall proton pumping efficiency of the respiratory chain, which may have increased reliance on glycolysis as an energy source [88–90]. Together, these metabolic changes suggest that *C. burnetii* both expanded its capacity to use host-derived carbon sources and refined control over central carbon metabolism, consistent with adaptation to a chemically complex intracellular compartment.

Several limitations should guide interpretation of these results. Comparative genomics can identify plausible gains, losses, duplications, and horizontal transfers, but targeted experiments will be needed to determine how these evolutionary changes affect *C. burnetii* physiology, intracellular growth, and virulence. For instance, some functional assignments, including parts of the virenose pathway, the mannose catabolism route, and the role of Mrp, require biochemical or genetic validation. Inferred evolutionary timing also depends on taxon sampling, orthology detection, and the availability of high-quality genomes from close relatives. Additional *Coxiella*-like bacteria, especially from understudied hosts, may refine the placement of individual acquisitions and losses. Even with these caveats, the major pattern is consistent across multiple systems: traits relevant to *C. burnetii* pathogenesis have different evolutionary depths and were combined gradually within a host-associated lineage, with ancestral functions later co-opted for survival and growth in the modern CCV.

Overall, our findings refine the evolutionary narrative of *C. burnetii*. Rather than emerging through abrupt acquisition of pathogenicity, *C. burnetii* is better understood as the product of a longer process in which ancestral host-interaction machinery was retained while envelope structure, stress tolerance, and metabolism were repeatedly modified. This framework helps explain how a member of Legionellales came to specialize in a vertebrate-associated, acidic intracellular niche and identifies specific evolutionary steps that can now be tested experimentally for their contributions to *C. burnetii* virulence and intracellular growth. More broadly, our results suggest that specialized intracellular pathogens can emerge through long-term remodeling of ancestral host-associated lineages, with older traits retained, repurposed, or combined with later gains and losses.

## MATERIALS AND METHODS

### Genome Sequencing and Assembly

A female *O. peruvianus* tick was collected from a cave inhabited by *Desmodus rotundus* bats on Pan de Azucar Island. DNA was extracted from the tick using the DNeasy Blood & Tissue Kit (Qiagen) and submitted to the Yale Center for Genome Analysis for Illumina (NovaSeq) sequencing. The resulting 150-bp paired-end reads were trimmed with Trimmomatic [91] using the following parameters: ILLUMINACLIP:2:30:10 SLIDINGWINDOW:5:25 LEADING:20 TRAILING:20 MINLEN:50. After trimming, approximately 490 million read pairs were retained. Reads were assembled into contigs with SPAdes v3.13.0 [92] using default parameters. Open reading frames were identified with Prodigal [93]. Contigs were then binned with CONCOCT [94] on the basis of coverage and k-mer composition. The *Coxiella*-containing bin was identified by BLASTn and BLASTp using a database of Coxiellaceae genomes. Approximately 13 million read pairs mapped to the *Coxiella*-containing bin and were used for the final assembly, yielding 112 contigs. Read mapping was performed with Bowtie2 [95] using a database of Coxiellaceae genome sequences. The *O. peruvianus* CE genome contains 106 highly conserved single-copy genes as defined by Albertsen et al. (2013) [96], comparable to the number typically found in *C. burnetii* strains. Final genome annotation was performed with the NCBI Prokaryotic Genome Annotation Pipeline [97]. Sequence data have been deposited under BioProject PRJNA1189154, BioSample SAMN44862134, SRA accession SRR31639864, and WGS master accession JBLIZD000000000. Assembly and annotation statistics for the *O. peruvianus* CE are provided in **Table 1**.

### Orthogroups

OrthoFinder v3.1.0 [98] was used to infer protein orthogroups for both tree construction and ancestral state reconstruction. The program was run multiple times with MCL inflation parameters of 1.2, 2.0, 2.5, and 3.0 to assess whether orthogroup assignments for focal genes were robust to clustering granularity. Orthogroup counts reported in **Table 1** were generated using an inflation parameter of 2.0. Protein-coding genes annotated as pseudogenized were translated into amino acid sequences and included in the orthogroup analysis. The full genome set used for orthogroup inference is listed in **Table S1**.

### Phylogenomic Analyses

The *Coxiella* genus-level phylogenomic tree was constructed from protein sequences encoded by 78 genes (**Table S2**), whereas the Legionellales tree was constructed from 152 genes (**Table S3**). In both cases, single-copy genes were identified from the orthogroup results, and genes annotated as pseudogenized were excluded from tree building. Individual protein sequences were aligned with MAFFT v7.490 [99] using the L-INS-i/globalpair iterative refinement strategy and then trimmed with trimAL v1.5.1 using the gappyout method [100]. The resulting alignments were concatenated and partition boundaries were recorded. Maximum-likelihood trees were inferred with IQ-TREE v3.0.1 [101]. ModelFinder was used for model selection, with the candidate set restricted to amino acid substitution matrices appropriate for bacteria. Branch support was estimated using 100 nonparametric bootstrap replicates.

Because many CEs have highly reduced genomes, including them in the order-level phylogenomic analysis substantially reduced the number of single-copy protein-coding genes available for tree construction. We therefore analyzed the *Coxiella* genus tree separately from the broader Legionellales tree to maintain robust gene sampling in both datasets. Coxiellaceae MAGs lacking species-level classification and originating from multi-isolate projects were evaluated for completeness before inclusion. MAGs without annotations were first annotated with Prokka v1.14.5 [102]. Genomes were excluded if they contained fewer than 100 of the highly conserved single-copy genes defined by Albertsen et al. (2013) [96], a threshold that excluded more than half of the available genomes. After preliminary tree construction, additional genomes were removed if their placements were unstable across analyses using different gene sets (70 to 152 genes) or if they consistently showed low branch support. The genomes retained in the final analyses, together with their accession numbers and host/source information are provided in **Table S1**.

Ancestral reconstruction and HGT inference were performed by applying Orthofinder results and the Legionellales and *Coxiella* ML trees (**Figure 1**) to PastML using the MPPA and F81 settings [98,103]. Pseudogenized orthologs were coded as present for ancestral-state reconstruction but interpreted as evidence of subsequent loss of function in extant taxa. For each *C. burnetii* gene, the origin node was determined to be the oldest node at which posterior probability was ≥0.7, with all subsequent nodes also having posterior probability of ≥0.7, as described previously [104,105]. Metabolic pathways and supplemental annotations were collected from KEGG using BlastKOALA [106,107], RAST Subsystems [108], and the Conserved Domain Database using RPS-BLAST [109]. Findings for focal genes were further confirmed using individual protein trees as described below.

For individual protein trees, homologs from taxa outside Legionellales were retrieved from the NCBI nonredundant protein (nr) database using BLASTp. The top 50 to 70 hits ranked by E-value were retained; the exact number varied when multiple hits shared the same E-value so that equivalent matches were not excluded arbitrarily. No more than five representatives from the same genus and no more than one representative from the same species were included. Alignments and maximum-likelihood trees were generated as described above. Branch support was evaluated with 1,000 ultrafast bootstrap replicates. Figures were prepared with iTOL v7.5.1 [110].

## DATA AVAILABILITY

Sequence data have been deposited under accessions JBLIZD000000000 (WGS master), PRJNA1189154 (BioProject), SAMN44862134 (BioSample), and SRR31639864 (SRA).

## ACKNOWLEDGMENTS

We thank Sebastian Munoz-Leal and Daniel Gonzalez-Acuna for collecting *Ornithodoros peruvianus* ticks. This work was supported in part by the University of Texas at San Antonio and National Institute of Allergy and Infectious Diseases grant AI126385.

## SUPPLEMENTAL INFORMATION

**Table S1.** Taxa used in analysis.

**Table S2.** Proteins used to build *Coxiella* phylogenomic tree.

**Table S3.** Proteins used to build Legionellales phylogenomic tree.

